# Magnetogenetic control of endogenous calcium signaling via ROS-sensitive ion channels reveals astrocytic regulation of neural circuit activity

**DOI:** 10.64898/2026.04.30.721963

**Authors:** Miriam Hernández-Morales, Juan Morales-Zúñiga, Tiffany Tran, Victor Han, Nikki Tjahjono, Ryan G Natan, Yuxiang Li, Lin Tian, Na Ji, Chunlei Liu

## Abstract

Astrocytes exhibit complex calcium signaling proposed to detect, integrate, and modulate neuronal activity; however, their causal contribution to neural circuit function remains unresolved. This is partly due to the lack of tools that recapitulate physiological astrocyte calcium signaling. Here, we show that experimental magnetic control of endogenous calcium signaling in astrocytes is sufficient to modulate neural circuit dynamics. Using pmFeRIC, a magnetogenetic tool for remote, noninvasive, and on-demand activation of endogenous calcium pathways, we demonstrate that selective activation of astrocyte calcium signaling drives neuronal activity. pmFeRIC uses magnetic fields and endogenous ferritin to generate localized reactive oxygen species (ROS) at the plasma membrane, activating native ROS-sensitive ion channels, inducing calcium influx. In hippocampal circuits *in vitro*, magnetic activation of endogenous astrocytic calcium signaling produced robust, state-dependent changes in neuronal circuit activity, in part by increasing presynaptic release probability, postsynaptic burst firing, and glutamatergic transmission. These effects were mediated by transient receptor potential channels, amplified by intracellular calcium stores, and abolished when astrocyte calcium signaling was disrupted, establishing a causal role in neural circuit dynamics. *In vivo*, pmFeRIC activates endogenous calcium signaling in astrocytes with kinetics comparable to locomotion-induced activity, enabling control of astrocyte calcium dynamics that resemble physiological behavioral responses. Together, these findings establish that endogenous astrocytic calcium signaling is sufficient to modulate neural circuit function and introduce a broadly applicable platform for cell-specific, wireless, on-demand control of endogenous calcium signaling through magnetic activation of ROS-sensitive ion channels.

## Introduction

Astrocytes are increasingly recognized as regulators of neural circuits that shape behavior^1–5^. They monitor and modulate synaptic function in part through Ca^2+^-dependent mechanisms^6–13^. In response to changes in extracellular potassium, neurotransmitters, and neuromodulators, astrocytes exhibit intracellular Ca^2+^ transients linked to astrocyte-to-neuron signaling, including the release of ATP^14,15^, GABA^16^, glutamate^17–19^, and D-serine^20,21^, capable of influencing synaptic transmission. However, the causal mechanisms by which astrocytes regulate neural circuits remain poorly defined. Resolving this question will determine whether astrocytes merely modulate neuronal activity or actively drive neural circuit function, shaping brain processes.

Astrocyte Ca^2+^ signaling has been linked to sensory processing, brain-state transitions, and experience-dependent behaviors^6,7,12,13,22–25^. Genetically encoded Ca^2+^ indicators (GECIs), such as GCaMP, have revealed complex intracellular Ca^2+^ dynamics in astrocytes associated with different behavioral states, rather than uniform Ca^2+^ elevations^6–8,12,26–31^. Astrocyte Ca^2+^ activity is heterogeneous across spatial and temporal scales, ranging from localized microdomain events in fine processes to cell-wide transients that can propagate between astrocytes^8,26,27,29^. Astrocytes also display spontaneous and neuron-independent Ca^2+^ activity^32–34^. Combining GECIs with genetically encoded actuators, such as those employed in optogenetics and chemogenetics, has enabled simultaneous monitoring and experimental manipulation of astrocyte Ca^2+^ signaling, providing an initial framework for testing causal mechanisms of astrocytes in neural circuits shaping behavior^1,6,7,10,16,23–25,35^. However, these tools rely on synthetic or recombinant ion channels, receptors, or ion pumps that can interfere with astrocyte physiology, may not engage endogenous Ca^2+^ pathways, and do not reproduce the spatiotemporal dynamics of astrocyte Ca^2+^ signaling observed in intact circuits.

Chemogenetics has limited temporal resolution; Ca^2+^ increases typically begin 10-40 minutes after clozapine-N-oxide administration and can persist for hours^36–38^.

Chemogenetic activation of astrocytes using G-protein– coupled DREADDs produces large, sustained Ca^2+^ elevations followed by refractory periods^25,38,39^, has off-target effects^40–42^ and can induce long-term transcriptional changes^43^. Optogenetics increases astrocytic Ca^2+^ signaling with temporal resolution. However, it requires recombinant actuators, can produce off-target effects, and is invasive. The widely used ChR2 channel can induce acidosis^46^ and alter intra-and extracellular potassium levels^47^. Moreover, the need for optical access, cranial window implantation, and high-intensity laser stimulation can disturb astrocyte physiology and introduce experimental artifacts^48^. Other strategies probe the role of astrocyte Ca^2+^ signaling by suppressing it using extrusion systems such as the human plasma membrane Ca^2+^ ATPase (CalEx)^44,45^, which chronically lowers intracellular Ca^2+^. Therefore, although these approaches link astrocyte calcium signaling to neural circuit activity, they do not establish a temporally defined role or reveal the causal mechanisms by which astrocytes regulate neuronal circuits.

Magnetogenetics has emerged as an alternative for noninvasive control of cellular activity^49–51^. Approaches that couple ferritin to recombinant TRPV channels enable radiofrequency (RF) magnetic fields to induce cellular Ca^2+^ responses^49–51^. Here, using the magnetogenetic tool FeRIC (Ferritin iron Redistribution to Ion Channels)^50,52–54^, we demonstrated RF-induced Ca^2+^ responses in astrocytes expressing the recombinant ferritin-coupled TRPV4. However, this approach robustly perturbs basal astrocytic Ca^2+^ activity by expressing recombinant TRPV4 channels. To overcome these limitations and those of existing tools, we developed pmFeRIC (plasma membrane FeRIC), a magnetogenetic strategy for precise and cell-type-specific control of endogenous Ca^2+^-permeable channels in astrocytes. pmFeRIC reallocates endogenous ferritin to the plasma membrane via an Lck-targeting motif fused to a ferritin linker (domain 5 of kininogen). The RF interaction with membrane-tethered ferritin induces local production of reactive oxygen species (ROS)^52,55,56^, which can modulate nearby native ROS-sensitive ion channels, including TRP channels. In hippocampal neural cultures, RF stimulation induces Ca^2+^ increases in pmFeRIC-expressing astrocytes mediated by endogenous TRPV4 channels. This activation triggers astrocyte-to-neuron signaling, enhancing neuronal spiking activity by increasing glutamatergic transmission. *In vivo*, RF triggers Ca^2+^ responses in pmFeRIC-expressing astrocytes in the visual cortex with kinetics resembling those observed during locomotion. By controlling native ROS-sensitive ion channels, pmFeRIC enables the experimental recapitulation of physiologically relevant Ca^2+^ dynamics, providing a powerful platform for causal dissection of astrocyte contributions to neural circuit function.

## Results

### Radiofrequency (RF) magnetic fields induced Ca^2+^ responses in astrocytes expressing TRPV4^FeRIC^

We first tested the FeRIC method^50,52–54^ using the recombinant ferritin-coupled TRPV4^FeRIC^ to control astrocytic Ca^2+^ signaling, monitored by employing GCaMP6 (cytosolic or lck) (**Fig. 1a-c, i**). Astrocytic Ca^2+^ signaling, detected as Ca^2+^ events at the processes, was quantified using the MATLAB AQuA algorithm^57^.

**Fig. 1.**
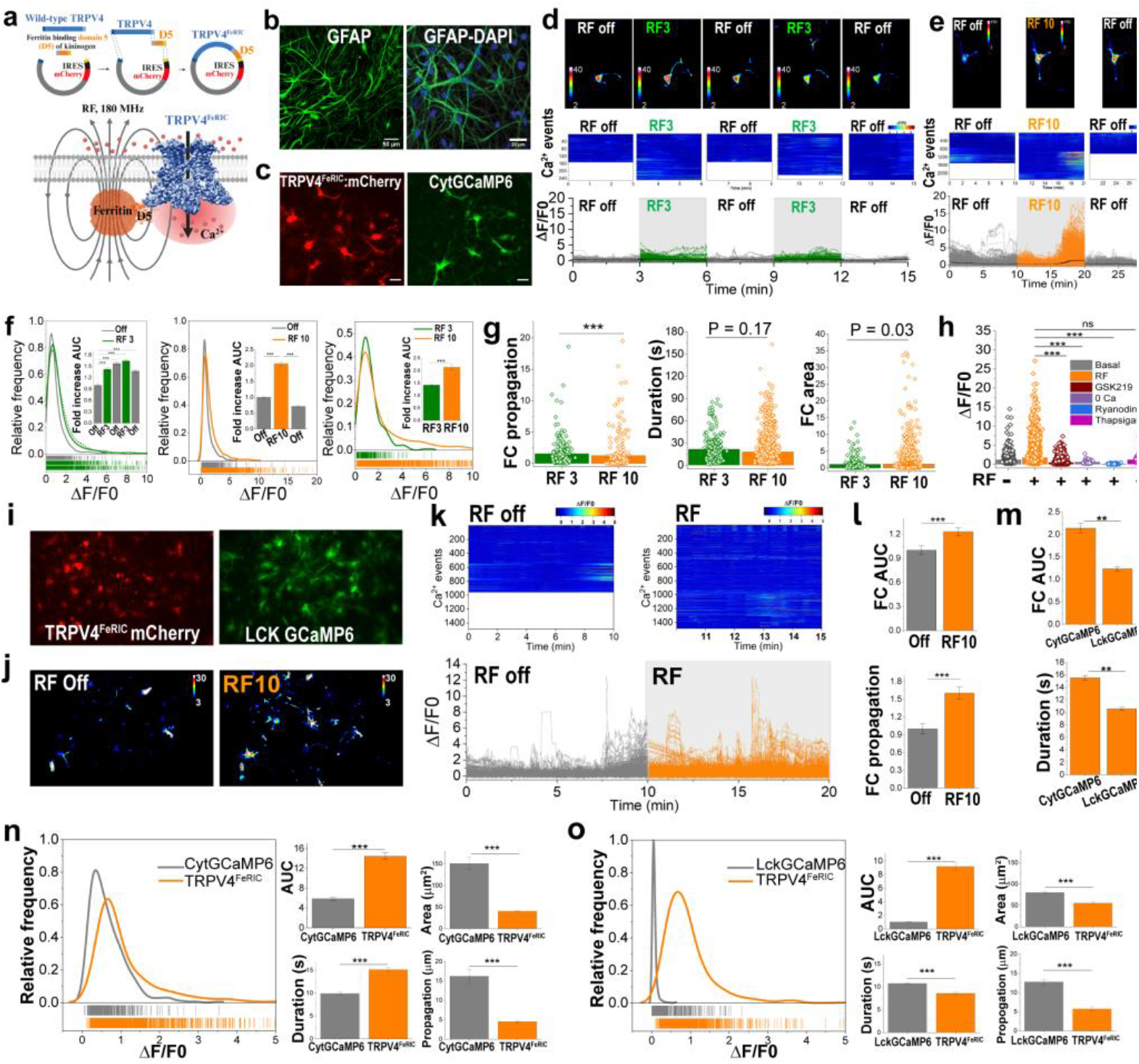
Magnetic control of astrocytic Ca^2+^ signaling using TRPV4^FeRIC^ channels. **(a)** Schematic of the FeRIC method. TRPV4^FeRIC^ channels were generated by fusing TRPV4 with the ferritin-binding domain 5 (D5) of kininogen. Endogenous ferritin binds D5, and radiofrequency (RF) fields activate TRPV4^FeRIC^ to induce Ca^2+^ influx. **(b)** Astrocyte-enriched cultures immunolabeled for GFAP and nuclei stained with DAPI. **(c)** Astrocytes co-expressing TRPV4^FeRIC^ (mCherry) and cytosolic GCaMP6. **(d – e)** Standard deviation maps (top), heat maps (middle), and cytGCaMP6 fluorescence traces (bottom) of Ca^2+^ events during RF stimulation for 3 min (d) or 10 min (e). **(f - g)** Distribution and mean (±SEM) of ΔF/F0 (F), propagation, duration, and area of Ca^2+^ events during RF off, or during 3-or 10-min RF stimulation. Insets: fold change of the AUC from cytGCaMP6 events without RF (off) or RF on. **(h)** Maximum cytGCaMP6 ΔF/F0 during TRPV4 blockade (GSK219), removal of extracellular Ca^2+^, or inhibition of SERCA and ryanodine receptors. **(i)** Astrocytes expressing TRPV4^FeRIC^ (mCherry) and the membrane-targeted v. **(j-k)** Standard deviation maps (j), heat maps, and fluorescence traces of lckGCaMP6 during 10-min RF off/on periods (k). **(l)** Mean (±SEM) of fold-change in the AUC and the propagation of Ca2+ events in astrocytes before (Off) and following RF exposure for 10 min **(m)** Comparison of fold change in AUC and duration when Ca^2+^ events are monitored with cytGCaMP6 versus lckGCaMP6. (**n-o**) Distribution of ΔF/F0, AUC, area, duration, and propagation observed in astrocytes expressing only (n) cyt-or (o) lckGCaMP6, or co-expressing these Ca^2+^ indicators with TRPV4^FeRIC^. For this and the following figures, except when indicated, data correspond to the averages from all FeRIC-or GCaMP6-expressing cells. For this and the following figures, except when indicated, significance was determined using the nonparametric two-sample Kolmogorov-Smirnov (two experimental groups) test or Kruskal-Wallis ANOVA followed by Dunn’s multiple comparisons test (three or more experimental groups). Where applicable, *P < 0.05, **P < 0.001, or ***P < 0.0001 was considered a statistically significant

We first tested FeRIC using astrocyte-enriched cultures. In control experiments with astrocytes not expressing TRPV4^FeRIC^, RF (180 MHz, 10 µT) did not increase GCaMP6 fluorescence (**Extended Data Fig. 1a, b**, cytGcaMP6: n = 4 independent experiments, 32 cells, 2020 – 4302 Ca^2+^ events; lckGCaMP6: n = 4 independent experiments, 19 cells, 1645 – 1714 Ca^2+^ events). In contrast, RF increased the number, amplitude, propagation, area, and duration of Ca^2+^ events in astrocytes expressing TRPV4^FeRIC^ (**Fig. 1c–m**; **extended Fig. 1c**; P < 0.0001, Kruskal–Wallis ANOVA with Dunn’s post hoc test). These effects were stimulation-duration dependent. Ten minutes of RF induced more frequent and larger Ca^2+^ events and increased the proportion of responsive cells compared with 3 min stimulation (AUC: 2.2 ± 0.05-fold, n = 2,805 events, 21 cells, 47.6% responsiveness versus AUC: 1.27 ± 0.06-fold, n = 248 events, 14 cells, 35.7% responsiveness) (**Fig. 1f**). RF effect on astrocytes was abolished by the TRPV4 antagonist GSK219, removal of extracellular Ca^2+^, and inhibition of ryanodine receptors or the SERCA pump (P < 0.0001, **Fig. 1h**). Similar results were observed when Ca^2+^ activity in astrocytes was monitored using lckGCaMP6 (P < 0.0001, **Fig. 1i-m, Extended data Fig.1 d-f**).

Although these results indicate that TRPV4^FeRIC^ enables magnetic manipulation of astrocytes, this channel induces aberrant basal increases in Ca^2+^ activity (**Fig. 1n,o**; P < 0.0001). Next, we assessed TRPV4^FeRIC^ subcellular localization and found it was predominantly retained in the ER, with ferritin detected as aggregates at the same location (**Extended Data Fig. 1g, h**). Thus, although TRPV4^FeRIC^ shows promise for controlling astrocytes, it presents notable limitations, including predominant retention in the ER and, more importantly, elevated basal Ca^2+^ activity in astrocytes. These features limit their suitability for probing the physiological roles of astrocytic Ca^2+^ signaling.

### pmFeRIC targets ROS-sensitive endogenous ion channels in astrocytes

To overcome TRPV4^FeRIC^ limitations, we developed the new FeRIC variant **pmFeRIC**, designed to activate endogenous ion channels in genetically targeted cells. pmFeRIC leverages RF interactions with ferritin to produce reactive oxygen species (ROS) to activate ROS-sensitive endogenous ion channels^52,55,56,58,59^. pmFeRIC consists of the membrane export lck signal fused with the ferritin-binding domain 5 (D5) from kininogen, resulting in a membrane-anchored ferritin linker (**Fig. 2a**). This new FeRIC variant enables selective, noninvasive control of endogenous intracellular Ca^2+^ signaling without recombinant ion channels, preserving intrinsic Ca^2+^ homeostasis and signaling dynamics (**Fig. 2b**).

**Fig. 2.**
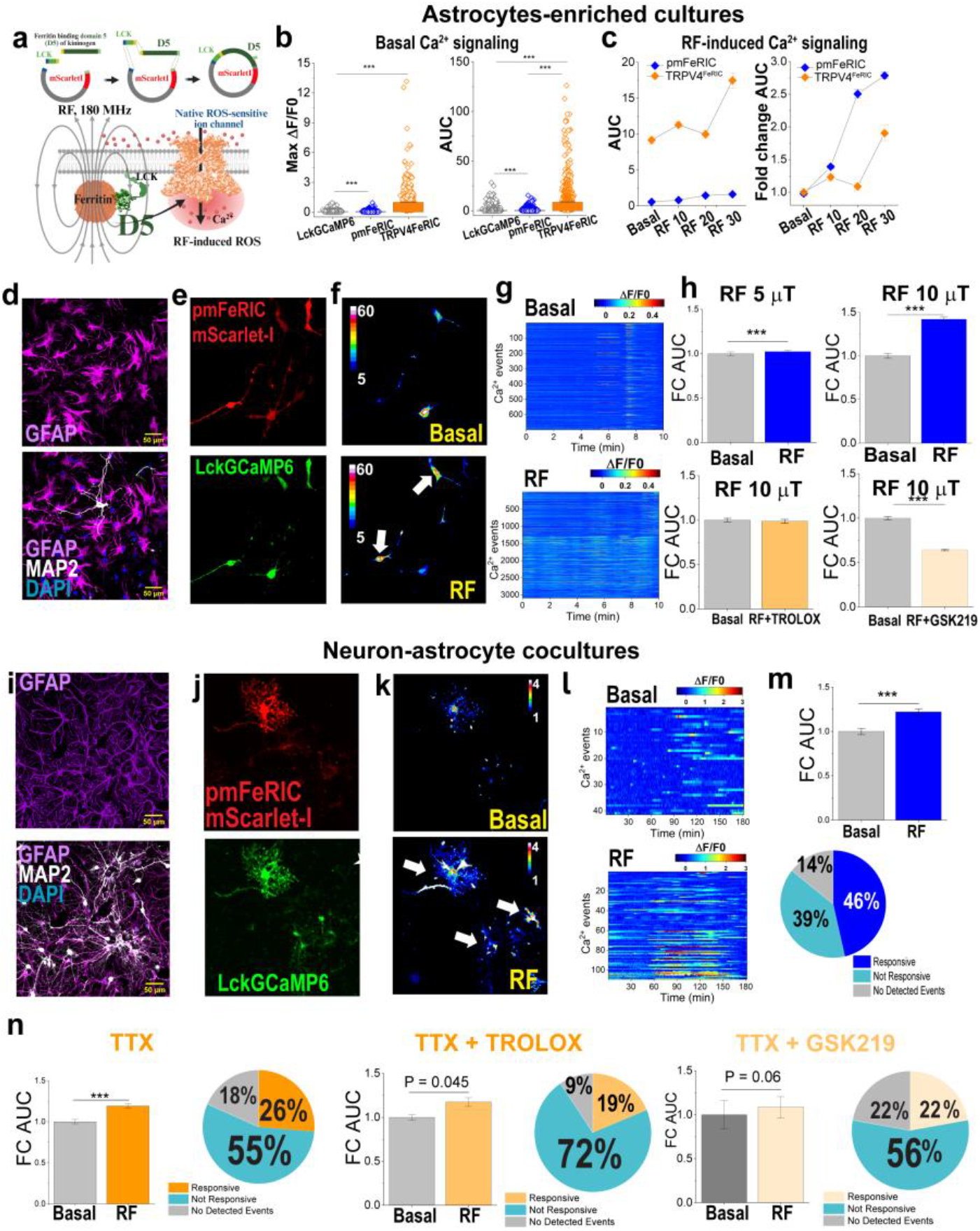
Magnetic activation of endogenous astrocytic ion channels induces Ca^2+^responses. (**a**) Schematic of the pmFeRIC variant. The ferritin-binding domain 5 (D5) is targeted to the plasma membrane via fusion to the Lck motif. Endogenous ferritin binds membrane-localized D5, and RF stimulation activates endogenous ion channels to induce Ca^2+^ influx. (**b**) Comparison of basal Ca^2+^ events amplitude (ΔF/F0 and AUC) in astrocytes expressing lckGCaMP6 alone, pmFeRIC, or TRPV4^FeRIC^. (c) Changes in lckGCaMP6 AUC and AUC fold change (mean ± SEM) during consecutive 10-min RF stimulations in astrocytes expressing pmFeRIC or TRPV4^FeRIC^. Fold change is normalized to the first 10-min baseline. (**d**) Enriched astrocyte cultures immunolabeled for GFAP and MAP2 (**e**) Astrocytes expressing pmFeRIC (mScarlet-I) and lckGCaMP6. (**f**,**g**) Pseudocolor SD images of Lck–GCaMP6 fluorescence (f) and corresponding Ca^2+^ event heat maps (g) in pmFeRIC-expressing astrocytes under basal and RF conditions (10 μT, 10 min). White arrows indicate astrocytes with increased intracellular Ca^2+^ levels. (**h**) Mean (± SEM) fold change in AUC in pmFeRIC-expressing astrocytes stimulated with 5 or 10 µT RF, or 10 µT RF in the presence of the TRPV4 antagonist GSK219 or the ROS scavenger Trolox. (**i**) Neuron–astrocyte co-cultures immunolabeled for GFAP and MAP2. (j) Astrocytes from co-cultures infected with AAV expressing pmFeRIC (mScarletI) and lckGCaMP6. (**k**,**l**) Pseudocolor images of lckGCaMP6 fluorescence (k) and corresponding Ca^2+^ event heat maps (l) during baseline and RF stimulation. (**m**) Fold change lckGCaMP6 AUC following 10-min RF stimulation at 10 µT and the percentage of RF-responsive astrocytes. (**n**) Fold change in lck:GCaMP6 AUC and the percentage of RF-responsive astrocytes following 10-min RF stimulation in the presence of TTX, TTX plus Trolox, or TTX plus GSK219. For Ca^2+^ imaging data, statistical analysis was performed as indicated in Fig.1.

In astrocyte-enriched cultures, we first confirmed pmFeRIC localization at the plasma membrane with no detectable ER retention of channels (**Extended Data Fig. 2a**), and that pmFeRIC does not alter basal astrocytic Ca^2+^ events. Astrocytes expressing pmFeRIC exhibited only a ∼1.02-fold increase in basal max ΔF/F0, in sharp contrast to the ∼17.8-fold increase observed in TRPV4^FeRIC^-expressing astrocytes relative to lck-GCaMP6 controls (lckGCaMP6 = 0.055 ± 0.0004-fold increase, pmFeRIC = 0.056 ± 0.0005-fold increase, TRPV4^FeRIC^: 0.98 ± 0.02-fold increase; P < 0.0001; independent experiments 4 – 6, cells 19 – 32; C Fig. 2b). Next, we confirmed membrane-localized pmFeRIC expression and verified the structural and functional presence of endogenous ROS-sensitive ion channels, including TRPA1, TRPV1, and TRPV4 (**Extended Data Fig. 2c**). We then compared RF-induced Ca^2+^ responses in astrocytes co-expressing pmFeRIC and lckGCaMP6 with those expressing TRPV4^FeRIC^. RF stimulation (10-min periods at 180 MHz and 10 µT) increased Ca^2+^ activity in both groups, with larger Ca^2^ event amplitudes in TRPV4^FeRIC^-expressing astrocytes. To enable direct comparison, we estimated the fold-change amplitude of Ca^2+^ events (**Fig. 2c**). In pmFeRIC-expressing astrocytes, consecutive 10-min RF stimulations produced 1.2-, 1.7-, and 1.8-fold increases in ΔF/F0. In TRPV4^FeRIC^-expressing astrocytes, the corresponding increases were 1.08-, 1.04-, and 1.4-fold. The difference was more evident in the AUC measurements: pmFeRIC-mediated responses increased from 1.4-to 2.8-fold, whereas those mediated by TRPV4^FeRIC^ responses increased from 1.08-to 1.9-fold. These results indicate that although TRPV4^FeRIC^-expressing astrocytes display larger amplitude Ca^2+^ events, the fold-change and the percentage of responsive cells are smaller compared with astrocytes expressing pmFeRIC (**Fig. 2c**). **Fig. 2d-h** shows detailed illustrations of RF-induced Ca^2+^ responses in astrocytes expressing pmFeRIC (AUC RF 5 µT = 1.02-fold increase, 6 independent experiments, 57 cells, 1172 – 1628 Ca^2+^ events; AUC RF 10 µT = 1.4-fold increase, 6 independent experiments, 32 cells, 5530 – 5769 Ca^2+^ events; P < 0.0001). RF-induced Ca^2+^ responses were dose-dependent, with 10 µT RF producing a larger fold-change in AUC (**Fig. 2h**) and a higher percentage of responsive cells compared with 5 µT RF (45% at 5 µT vs. 53% at 10 µT). However, when cells were segregated based on their responsiveness, the RF-responsive population showed similar fold-changes in AUC at 5 and 10 µT (**Extended Data Fig. 2d,e**). These results suggest that increasing RF strength recruits more responsive cells rather than enhancing the amplitude of Ca^2+^ events. Next, we verified that RF activates ROS-sensitive TRPV4 channels because Trolox and GSK219 abolished RF effects (Trolox: AUC: 0.99-fold change FC; 3 independent experiments, 8 cells, 1885 – 3425 Ca^2+^ events; GSK219 AUC: 0.6-FC, 2 independent experiments, 18 cells, 8562 - 8580 Ca^2+^ events; P < 0.0001; **Fig. 2h**).

We corroborated similar results in neuron-astrocyte cocultures (**Fig. 2i**, see Methods), an *in vitro* model suitable for assessing Ca^2+^-dependent astrocytic modulation of neuronal activity. To selectively express pmFeRIC and GCaMP6 in astrocytes, we used an AAV delivery system with the astrocyte-specific gfaABC1D promoter (**Fig. 2j**). We confirmed the expression of endogenous TRP channels in astrocytes from cocultures (**Extended Fig. 2f**). Next, we corroborated that pmFeRIC activates endogenous channels mediating Ca^2+^ responses in ∼46% of astrocytes upon RF exposure (AUC: 1.22-fold increase; 6 independent experiments; 28 cells; 1,172 - 1,628 Ca^2+^ events) (**Fig. 2j–m**). To assess dependence on neuronal activity, we applied the Na^+^ channel blocker tetrodotoxin (TTX). Under TTX, RF induced a similar AUC fold-change (AUC: 1.19-fold increase; 6 independent experiments; 38 cells; 2,101 – 2,222 Ca^2+^ events), but the percentage of responsive cells was reduced to 26% (**Fig. 2j,n**). Trolox (AUC: 1.17-fold; 6 independent experiments; 32 cells; 596 – 672 Ca^2+^ events, 26 % of RF responsive cells) or GSK219 (AUC: 1.08-fold increase; 3 independent experiments; 9 cells; 65 – 85 Ca^2+^ events, 22 % of RF responsive cells) further decreased the proportion of responsive cells, the AUC amplitude (**Fig. 2n**), and the number of Ca^2+^ events per cell. These results indicate that pmFeRIC drives astrocytic Ca^2+^ responses in *in vitro* neural networks by activating endogenous ROS-sensitive TRPV4 channels.

To further examine pmFeRIC targeting native membrane ion channels, we used whole-cell voltage-clamp recordings. Astrocytes expressing pmFeRIC were identified by mScarlet-I fluorescence, and membrane currents were recorded under sham or RF stimulation (180 MHz, 10 µT). Astrocytes exhibited both inward and outward currents (**Fig. 3a,b**); because outward currents were sporadic, subsequent analyses focused on inward currents. Astrocytes displayed a diversity of inward currents, which were grouped based on amplitude and kinetics: Type I (>10 pA) with slow rise (several seconds) and decay (minutes); Type II (>10 pA) with activation within ∼3 s and decay over hundreds of seconds; Type III (>10 pA) with fast activation and decay (∼1–3 s); and Type IV, small-amplitude currents (<10 pA) with the fastest kinetics.

**Fig. 3.**
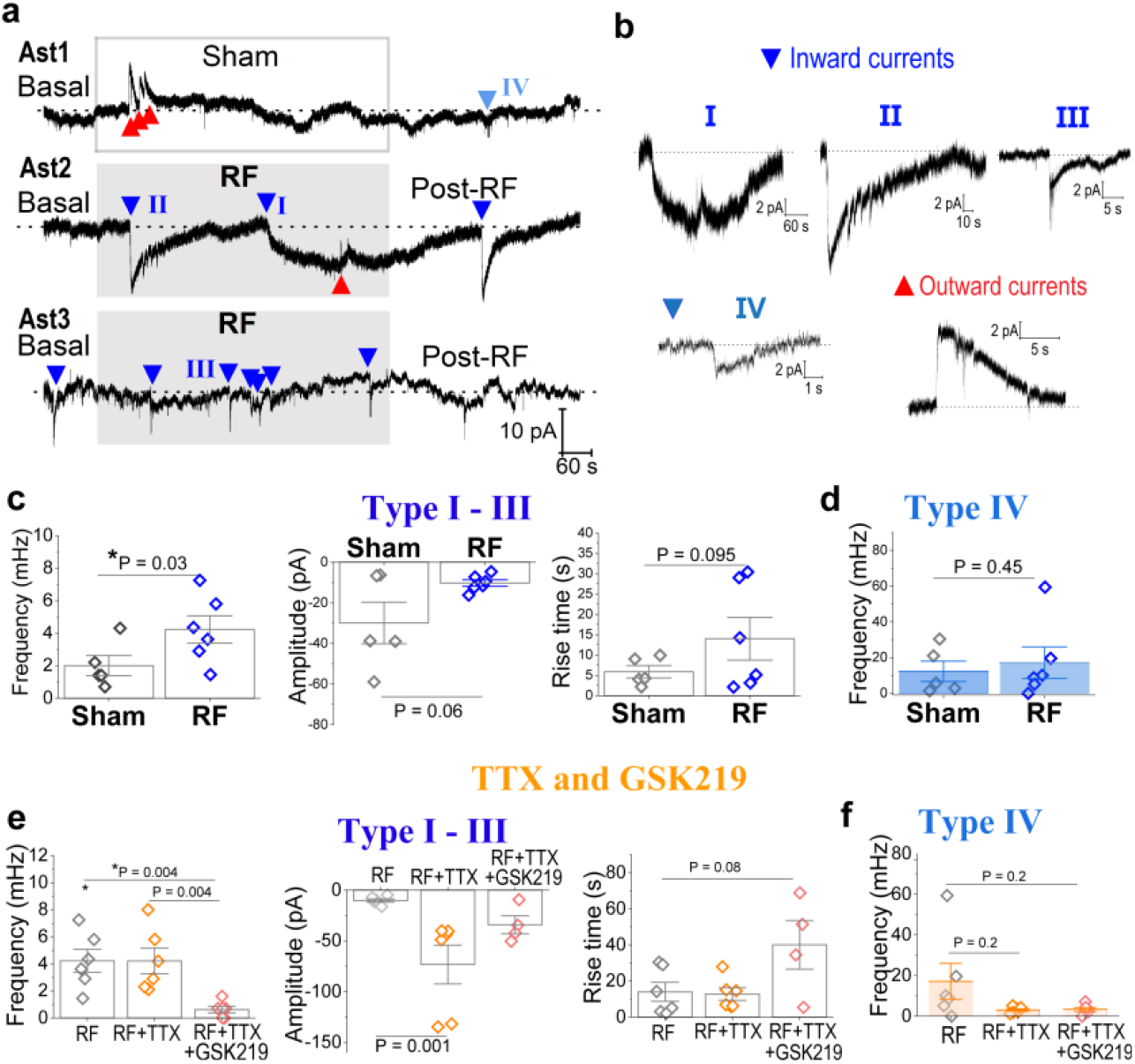
pmFeRIC targets endogenous membrane ion channels and enables RF-induced cation currents in astrocytes. (**a**) Representative voltage-clamp recordings from astrocytes during sham (top) or RF stimulation (middle, bottom). Red and blue arrows indicate outward and inward currents, respectively. (**b**) Inward currents were classified into four types (I–IV) based on amplitude, duration, and kinetics. Types I–III (>10 pA) were analyzed collectively. (**c**,**e**) Frequency, amplitude, and rise time of type I–III inward currents under sham (n = 5 cells) or RF stimulation (n = 6 cells; 10 µT, 10 min) in the absence (c) or presence of TTX (n = 6 cells) or TTX + GSK219 (n = 5 cells; e). (**d**,**f**) Frequency of type IV inward currents under sham or RF stimulation in the absence (d) or presence of TTX or TTX + GSK219 (f). Significance was determined using the parametric Two-sample T-test, and P values are indicated.

Under sham conditions, pmFeRIC-expressing astrocytes predominantly exhibited Type IV currents (**Fig. 3a**). RF significantly increased the frequency of large-amplitude inward currents (Types I–III) compared with sham (Sham: 2.0 ± 0.6 mHz, n = 5 cells; RF: 4.3 ± 0.8 mHz, n = 6 cells; P = 0.03), without significantly altering current amplitude (Sham: -30 ± 10.2 pA; RF: -10 ± 1.65 pA) or rise time (Sham: 5.86 ± 1.5 s; RF: 14 ± 5.2 s; **Fig. 3c**). In contrast, RF stimulation did not affect the frequency of Type IV events (Sham: 12.3 ± 5.7 mHz; RF: 17.2 ± 8.8 mHz; P = 0.45, **Fig. 3d**). To assess whether RF-induced currents depend on neuronal activity, recordings were performed in the presence of TTX, which did not alter the frequency of RF-evoked large inward currents (Types I–III: 4.2 ± 0.9 mHz; P = 1, n = 6 cells), but reduced the frequency of small Type IV currents, although this effect did not reach significance (3.0 ± 0.6 mHz; P = 0.2, n = 6 cells; **Fig. 3e,f**).

To examine the contribution of TRPV4 channels, astrocytes were treated with TTX and the TRPV4 antagonist GSK219. This treatment reduced large inward currents (Types I–III: 0.6 ± 0.2 mHz; P = 0.004, n = 6 cells), whereas the frequency of Type IV events did not change (3.3 ± 1.0 mHz; P = 1, n = 6 cells) (**Fig. 3e,f**). Together, these results corroborate that pmFeRIC enables RF to induce inward cation currents in astrocytes independently of neuronal activity. Our data suggest that the large-amplitude inward currents are RF-dependent and mediated largely by TRPV4 channels, whereas small-amplitude Type IV events are insensitive to RF and reduced by TTX. The diversity of inward current kinetics suggests the involvement of multiple endogenous TRP channels, with TRPV4 as the main contributor.

### Magnetic activation of endogenous astrocytic ion channels modulates neuronal activity via presynaptic signaling

To investigate the impact of RF-induced astrocytic Ca^2+^ signaling on neuronal activity, we performed Ca^2+^ pmFeRIC under the gfaABC1D promoter and neurons expressed GCaMP6f under the hSyn promoter (**Fig. 4a**). This configuration enabled selective RF activation of astrocytes while monitoring neuronal Ca^2+^ activity. Neurons exhibited fast Ca^2+^ transients consistent with action potential firing. We quantify changes in neuronal Ca^2+^ spiking frequency with a customized Python algorithm (see Analysis). We estimated the Ca^2+^ spiking frequency, and all neurons were included in the Analysis to capture the population-level effect. In our conditions, neurons exhibited heterogeneous Ca^2+^ spiking profiles: (1) increased spiking, (2) slow Ca^2+^ elevations with minimal spiking, and (3) slow Ca^2+^ elevations with superimposed spikes (**Fig. 4b, extended Fig. C**). We first established that RF does not directly affect neuronal Ca^2+^ spiking and that under basal conditions, Ca^2+^ spiking frequency did not differ between cocultures lacking and those containing pmFeRIC-expressing astrocytes (P = 1). In neurons expressing only hSyn-GCaMP6f, neither prolonged imaging (up to 30 min, 3 independent experiments, 55 cells, P = 0.2 to 1) nor RF stimulation increased neuronal Ca^2+^ spiking (5 independent experiments, 88 cells, P = 0.03; **Extended Data Fig. 4a**). Similarly, long-term imaging of cocultures containing pmFeRIC-expressing astrocytes did not change basal neuronal Ca^2+^ spiking in the absence of RF (10 min: 35 ± 3 mHz, 20 min: 43 ± 4 mHz, 30 min: 43 ± 3 mHz; 3 independent experiments, 70 cells **Fig. 4d**). In contrast, RF significantly increased neuronal Ca^2+^ spiking frequency in cocultures containing pmFeRIC-(basal: 20 ± 2 mHz, RF: 37 ± 2, post-RF: 31 ± 2 mHz: 15 independent experiments, 198 cells, P = 0.0035; **Fig. 4c,d Extended Data Fig. 4b**). RF-induced effects on neurons surrounded by pmFeRIC-astrocytes were abolished with the antioxidant Trolox (basal: 35 ± 4 mHz, RF: 41 ± 3, post-RF: 32 ± 4 mHz: 3 independent experiments, 28 cells, P = 0.33 to 0.79) and GSK219 (basal: 41 ± 3 mHz, RF: 40 ± 4, post-RF: 31 ± 2 mHz: 7 independent experiments, 108 cells, P = 0.20 to 1) (**Fig 4d, extended Fig. 4c**). To observe the astrocyte-to-neuron signaling independent of synaptic transmission, we imaged neurons from cocultures with pmFeRIC-astrocytes in the presence of TTX. Under these conditions, RF induced a robust increase in slow neuronal Ca^2+^ responses without spiking (AUC/s basal: -0.0085 ± 0.005, RF: 0.93 ± 0.08, post-RF: 1.14 ± 0.12, independent experiments 5, 54 cells) (**Fig. 4f; extended data Fig. 4d**). These responses were blocked by GSK219 (AUC/s RF+TTX+GSK219: 0.1 ± 0.14, 2 independent experiments, 43 cells) as well as by antagonists of NMDA, kainate, and AMPA receptors (AUC/s RF+TTX+2AP5: 0.46 ± 0.03, RF+TTX+CNQX: 0.26 ± 0.03, RF+TTX+CNQX: 0.35 ± 0.04, 2-4 independent experiments, 43 - 113 cells, P < 0.001, P < 0.0001) (**Fig. 4f,g; extended data Fig. 4e,f**).

**Fig. 4.**
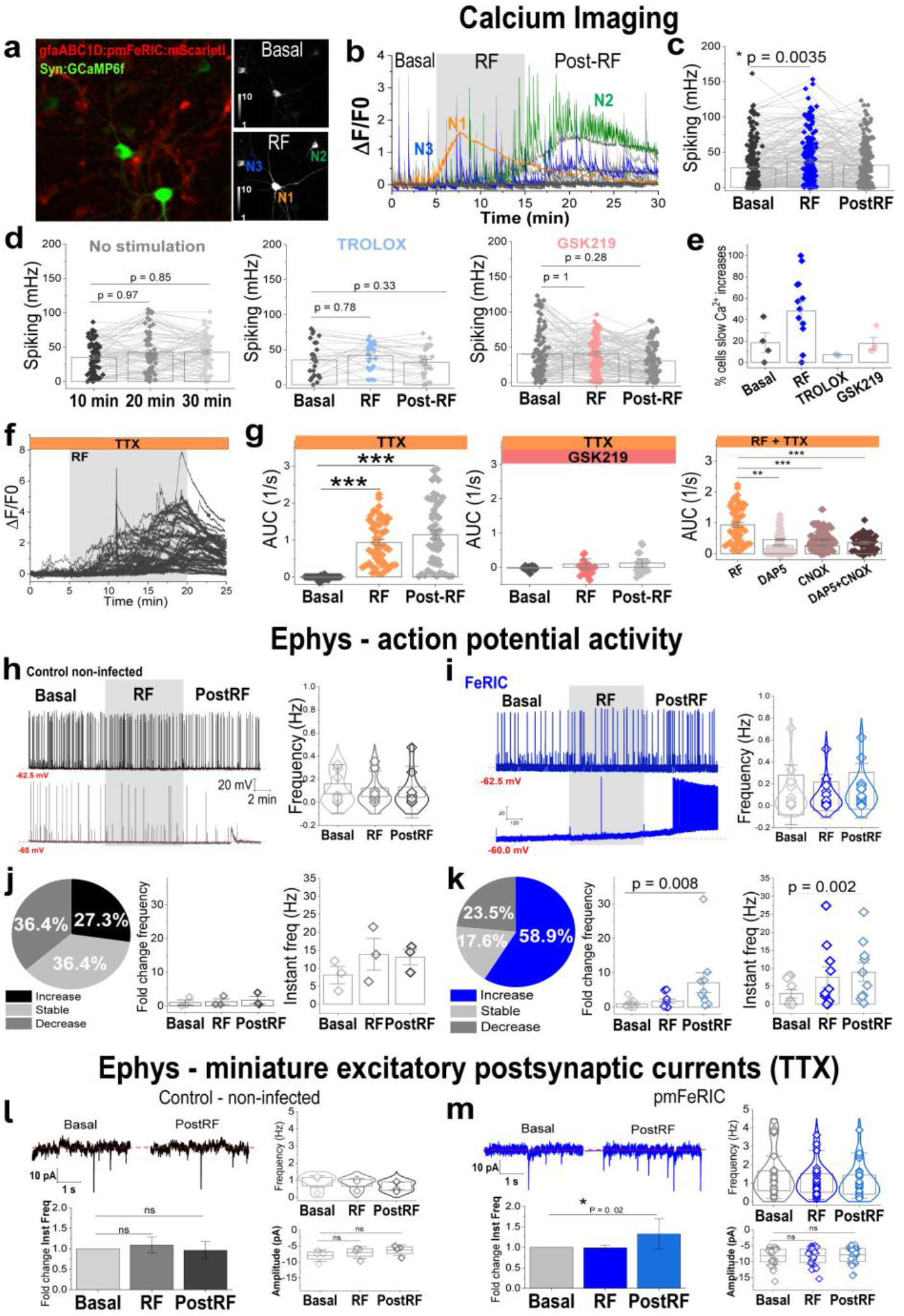
pmFeRIC astrocyte–mediated control of neuronal activity. (**a**) Neuron–astrocyte cocultures expressing hSyn:GCaMP6f in neurons and astrocyte-specific pmFeRIC. Right, standard deviation maps of hSyn:GCaMP6f fluorescence before (top) and after (bottom) RF stimulation. (**b**) Representative hSyn:GCaMP6f traces during basal, RF (gray bar), and post-RF periods, illustrating slow Ca^2+^ elevations (N1), mixed slow and spiking activity (N2), and spiking activity (N3), corresponding to color-coded neurons in (a). (**c**,**d**) Mean ± SEM neuronal Ca^2+^ spiking frequency during basal, RF, and post-RF periods in cocultures with pmFeRIC-expressing astrocytes under sham (3 independent experiments, 70 cells) or RF stimulation (10 µT; 15 independent experiments, 198 cells). (**f**,**g**) In separate experiments, RF-stimulated cultures were treated with the ROS scavenger Trolox (3 independent experiments, 28 cells) or the TRPV4 antagonist GSK219 (7 independent experiments, 108 cells). Each datapoint represents the average activity of individual neurons. (**e**) Fraction of neurons exhibiting slow Ca^2+^ increases during basal conditions, RF stimulation, or RF in the presence of ROS scavenging or TRPV4 blockade. To assess synaptic transmission contribution, cocultures were treated with the Na^+^ channel blocker TTX (1 µM). Representative hSyn:GCaMP6f traces (**f**) and area under the curve (AUC; mean ± SEM; **g**) during basal, RF, and post-RF periods in the presence of TTX alone (n = 4 independent experiments, 54 cells), TTX + GSK219 (n = 2 independent experiments, 43 cells), or TTX combined with ionotropic glutamate receptor antagonists (CNQX: n = 4 experiments, 113 cells; D-AP5: n = 4 experiments, 104 cells; CNQX + D-AP5: n = 2 independent experiments, 39 cells). (**h**,**i**) Representative current-clamp recordings from neurons in cocultures without (uninfected; h) or with pmFeRIC-expressing astrocytes (i) during basal, RF (10 µT; gray bar), and post-RF periods. Vertical traces are action potentials. Top traces show highly active neurons, whereas bottom traces show low-activity neurons. Bars are the mean action potential frequency (±SD) across conditions. (**j**,**k**) Proportion of neurons exhibiting increased, decreased, or unchanged firing frequency, and corresponding fold change and instantaneous firing frequency during RF stimulation in cocultures without or with pmFeRIC-expressing astrocytes (uninfected: n = 11 neurons; pmFeRIC: n = 17 neurons). (**l**,**m**) To assess synaptic transmission, neurons were recorded in the presence of TTX to measure miniature excitatory postsynaptic currents (mEPSCs) during basal, RF, and post-RF periods in cocultures without (l) or with pmFeRIC-expressing astrocytes (m). Representative mEPSC traces before and after RF stimulation (10 µT, 10 min) are shown. Summary data includes mEPSC frequency, fold change in instantaneous frequency, and amplitude (uninfected: n = 4 neurons; pmFeRIC: n = 29 neurons). For Ca^2+^ imaging data, statistical significance was assessed as indicated in Fig.1.

To further characterize astrocyte-mediated modulation of neuronal activity, we performed current-clamp recordings from neurons adjacent to astrocytes expressing pmFeRIC or lacking pmFeRIC expression (non-infected controls). Spontaneous action potential firing was recorded before, during, and after RF stimulation (**Fig. 4h,i**). Neurons exhibited heterogeneous average firing frequencies (0-0.7 Hz). All recorded neurons were included in the analysis to capture population-level effects. The proportion of neurons exhibiting increased firing rose from ∼27% in control cocultures to nearly 60% in cocultures containing pmFeRIC-expressing astrocytes upon RF stimulation (**Fig. 4j,k**). Although absolute firing rate showed only a trend toward increased activity in neurons adjacent to pmFeRIC-expressing astrocytes stimulated with RF, this did not reach statistical significance due to large inter-neuronal variability (uninfected: basal 0.12 ± 0.04 Hz, RF 0.08 ± 0.04 Hz, post-RF 0.08 ± 0.04 Hz, n = 11; pmFeRIC: basal 0.10 ± 0.04 Hz, RF 0.09 ± 0.03 Hz, post-RF 0.14 ± 0.04 Hz, n = 17) (**Fig. 4j,k**). To account for this variability, we quantified the fold change relative to baseline only in cells that increased the firing. This analysis revealed a significant increase in firing frequency in neurons adjacent to pmFeRIC-expressing astrocytes upon RF stimulation, with no change in control cultures (non-infected post-RF: 0.76 ± 0.4; pmFeRIC post-RF: 7.1 ± 1.4; P = 0.008; **Fig. 4j,k**). Consistently, RF significantly increased instantaneous firing frequency, reflecting changes in spike timing and bursting dynamics, in neurons neighboring pmFeRIC-expressing astrocytes but not in controls (P = 0.002; **Fig. 4j,k**). To assess whether these effects reflected changes in intrinsic neuronal excitability, we analyzed action potential threshold, resting membrane potential, and their difference. None of these parameters changed before, during, or after RF stimulation in any of these conditions (**Extended Data Fig. 4f,g**), indicating that astrocyte-mediated modulation does not involve postsynaptic changes in membrane excitability.

To determine whether astrocytes modulate neuronal activity presynaptically, we performed voltage-clamp recordings in neurons from cocultures treated with TTX. Miniature excitatory and inhibitory postsynaptic currents were observed, and analysis focused on mEPSCs. RF increased the proportion of neurons showing increased mEPSC frequency following RF stimulation in pmFeRIC cocultures (∼25% to ∼45%), although absolute frequencies were highly variable and did not differ significantly between conditions (non-infected, basal: 1.4 ± 0.2 Hz, RF: 1.1 ± 0.2 Hz, post-RF: 1.1 ± 0.2 Hz; pmFeRIC, basal: 1 ± 0.2 Hz, RF: 0.9 ± 0.2 Hz, post-RF: 0.85 ± 0.2 Hz) (**Fig. 4l,m**). In contrast, analysis of fold changes in instantaneous mEPSC frequency revealed a significant RF-induced increase in pmFeRIC cocultures (non-infected, RF: 1 ± 0.06, post-RF: 1 ± 0.2 Hz, n = 4 cells; pmFeRIC, RF: 1 ± 0.06 Hz, post-RF: 1.3 ± 0.4 Hz, n = 29, P = 0.02) with no changes in mEPSC amplitude or amplitude fold change (non-infected, basal: -8 ± 0.9 pA, RF: -7 ± 0.9 pA, post-RF: -6.3 ± 0.8 pA, n = 4 cells; pmFeRIC, basal: -8.2 ± 0.5 pA, RF: -8.1 ± 0.5 pA, post-RF: -7.9 ± 0.5 pA, n= 9 cells) (**Fig. 4l,m**). Consistently, peak-scaled non-stationary fluctuation analysis revealed no change in unitary current or number of postsynaptic glutamate receptors (**Extended Data Fig. 4h**), suggesting astrocytes do not affect neurons by postsynaptic mechanisms.

Together, these results indicate that RF activation of pmFeRIC-expressing astrocytes enhances neuronal firing by increasing presynaptic glutamatergic transmission, consistent with an elevated probability of neurotransmitter release. This astrocyte-driven modulation occurs without changes in postsynaptic excitability or synaptic receptor number.

### Astrocyte-specific magnetic activation in V1 cortex in vivo

Next, we examined whether pmFeRIC enables control of astrocytic Ca^2+^ signaling *in vivo*. Mice were injected unilaterally into the left primary visual cortex (V1) with AAV-pmFeRIC and AAV-cytGCaMP6, both driven by the astrocyte-specific gfaABC1D promoter. Astrocytic Ca^2+^ signaling was assessed by two-photon imaging in layer II/III of V1 in head-fixed mice under two conditions: anesthetized mice during RF stimulation and awake mice during spontaneous locomotion. Under anesthesia, mice do not locomote, ruling out movement as a contributor to RF-evoked Ca^2+^ responses. Awake imaging was performed specifically to capture locomotion-induced astrocytic Ca^2+^ activity, ensuring that similarities between RF-and locomotion-evoked responses are not attributable to animal motion. Expression of pmFeRIC and GCaMP6 was confirmed by mScarlet-I and GCaMP6 fluorescence (**Fig. 5a,b**), and mScarlet-I colocalized with the astrocytic marker GFAP at the ipsilateral injection site but not contralaterally (**Fig. 5c**). We first established that the head-sized RF coil did not affect astrocytic Ca^2+^ activity in anesthetized mice expressing GCaMP6 alone (1.1 ± 0.01-FC in ΔF/F_0_; 19 experiments, 3 mice; P = 0.2; Extended Data Fig. 5a,b) or in pmFeRIC-and GCaMP6-expressing anesthetized mice under sham conditions, in which the RF system was powered but no magnetic field was delivered (0.98-FC; 19 experiments, 6 mice; P = 0.5) (**Fig. 5d,e)**. As a positive control for astrocyte responsiveness, we imaged astrocytic Ca^2+^ activity in awake mice during spontaneous locomotion. Locomotion elicited Ca^2+^ elevations in ∼20% of astrocytes, with events lasting ∼8 s and temporally aligned across astrocytes in the FOV. All Ca^2+^ event parameters were significantly increased (1.3 ± 0.08-FC in ΔF/F0; 13 experiments, 5 mice; P < 0.0001; **Fig. 5f,g**). We then applied RF stimulation (50 µT) for 1, 5, or 10-min in anesthetized mice. RF induced rapid astrocytic Ca^2+^ elevations in the anesthetized brain that closely resembled locomotion-evoked responses in awake mice (**Fig. 5h-j; Extended Data Fig. 5c,d**). Across stimulation durations, RF elicited Ca^2+^ responses with comparable amplitudes and durations (1 min: 1.3 ± 0.2, 8.5 ± 0.9 s; 5 min: 1.2 ± 0.04, 10.2 ± 0.6 s; 10 min: 1.25 ± 0.04, 8.0 ± 0.2 s; 3–21 experiments, 4 mice; P = 0.3–1; **Fig. 5k**), but longer RF stimulation increased the percent of RF responsive cells (**Fig. 5l**). Repeated RF stimulation in the same mouse (three trials separated by 20 min) produced nearly identical Ca^2+^ responses, demonstrating reproducibility (**Extended Data Fig. 5c**). Similar responses were also observed across imaging sessions on different days. Although RF-induced Ca^2+^ responses were comparable when quantified as fold change, their absolute amplitude scaled with mScarlet-I fluorescence, a proxy for pmFeRIC expression level. mScarlet-I intensity correlated with maximal ΔF/F0 and fold change, whereas GCaMP6 expression did not, indicating that response magnitude reflects pmFeRIC rather than GCaMP6 expression (**Extended Data Fig. 5e**). Finally, we compared RF-and locomotion-induced Ca^2+^ responses. Both conditions produced similar amplitudes and durations, but differed in spatial features such as event area and propagation (**Extended Data Fig. 5f**). This distinction is consistent with locomotion engaging neuromodulatory pathways, whereas RF directly activates astrocytes via pmFeRIC targeting endogenous ion channels. The similarity in response kinetics and magnitude supports the physiological relevance of RF-evoked astrocytic Ca^2+^ signaling *in vivo*.

**Fig. 5.**
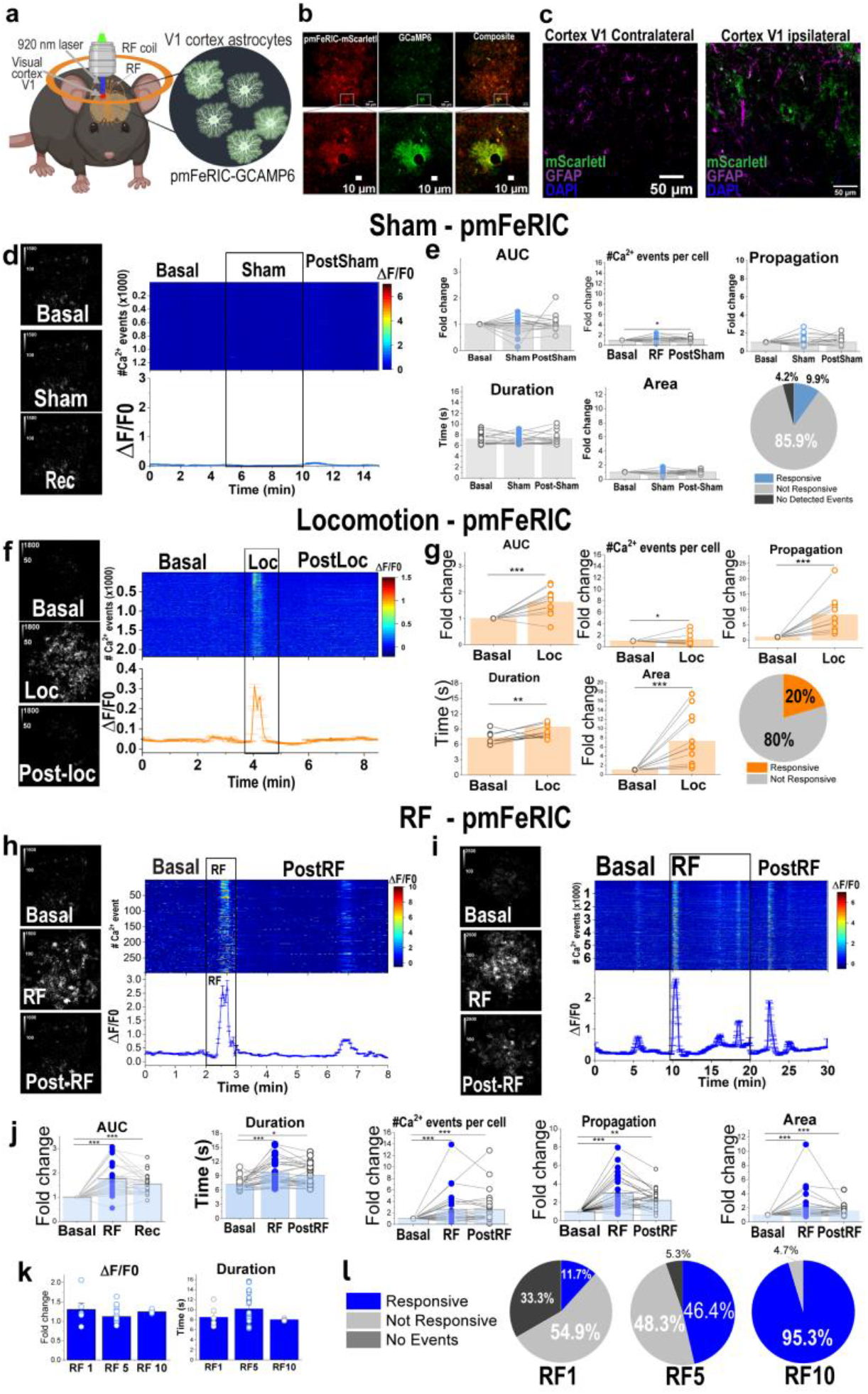
Magnetic activation of endogenous ion channels in pmFeRIC-expressing astrocytes in vivo. (**a**) Schematic of in vivo two-photon imaging in the visual cortex (V1) of a mouse expressing pmFeRIC and cytGCaMP6 in astrocytes. (**b**) Representative field of view from V1 cortex with magnified images showing astrocytes expressing pmFeRIC (mScarletI) and cytGCaMP6. (**c**) Contralateral (no injected) and ipsilateral (injected) V1 brain sections immunolabeled to detect pmFeRIC (mScarletI, green) and the astrocyte marker GFAP (magenta). Nuclei are stained with DAPI. (**d**,**f**,**h**,**i**) Representative cytGCaMP6 fluorescence images, their corresponding Ca^2+^ event heat maps, and cytGCaMP6 traces from pmFeRIC-expressing astrocytes in V1 cortex during sham, locomotion, or RF stimulation (50 µT for 1 or 10 min). (**e**,**g**,**j**) Quantification of cytGCaMP6 AUC fold change, number of Ca^2+^ events, event propagation, event duration, and event area (mean ± SEM), as well as the percentage of responsive astrocytes, during sham (4 mice, 22 independent experiments, 1116 cells), locomotion (3 mice, 13 independent experiments, 205 cells), or RF stimulation for 1 (2 mice, 6 independent experiments, 264 cells) or 10 min (2 mice, 3 independent experiments, 86 cells). (**k**) Comparison of Ca^2+^ event amplitude (ΔF/F0) and duration in V1 astrocytes expressing pmFeRIC following 1-, 5-, or 10-min RF stimulation, and (**i**) proportion of responsive astrocytes for each RF stimulation duration. For 5 min RF stimulation data corresponds to 3 mice, 26 independent experiments, 1300 cells. Statistical significance was assessed as indicated in Fig.1, with the following P values: *P < 0.05, **P < 0.001, or ***P < 0.0001.

## Discussion

Here we introduce pmFeRIC (plasma membrane Ferritin-iron Redistribution to Ion Channels), a genetically encoded method for cell-type-specific activation of endogenous ion channels using radiofrequency (RF) magnetic fields. We show that pmFeRIC enables noninvasive, on-demand control of astrocytic Ca^2+^ activity by activating endogenous ROS-sensitive TRP channels, inducing intracellular Ca^2+^ elevations that mediate astrocytic modulation of neuronal activity. By avoiding the expression of recombinant receptors or channels, pmFeRIC preserves physiological astrocytic Ca^2+^ dynamics and, by targeting endogenous ion channels, more closely recapitulates native astrocyte signaling. This feature distinguishes pmFeRIC from existing neuromodulation tools and enables the interrogation of astrocytic signaling with physiological fidelity.

Our initial FeRIC approach uses RF activation of recombinant TRPV4 channels coupled to endogenous ferritins to control cellular activity^50,53^. Although TRPV4^FeRIC^ mediates RF-induced Ca^2+^ responses in astrocytes, its expression altered basal astrocytic Ca^2+^ dynamics, increasing the amplitude and duration of spontaneous events while restricting their spatial spread. Such alterations preclude physiological interrogation of astrocytic Ca^2+^ signaling and mirror limitations of other approaches, such as chemogenetics, which often induce sustained, non-physiological Ca^2+^ elevations and prolonged refractory periods^60,61^. In contrast, pmFeRIC targets endogenous, ROS-sensitive ion channels, preserving basal Ca^2+^ dynamics and enabling physiologically relevant manipulation of astrocytic signaling for defining causal astrocyte–neuron interactions in neural circuits.

pmFeRIC operates by tethering endogenous ferritin to the plasma membrane to induce *in situ* ROS production upon RF exposure thereby activating ROS-sensitive membrane ion channels. Using Ca^2+^ imaging and patch-clamp recordings, we demonstrate that RF activates endogenous plasma membrane ion channels inducing astrocyte Ca^2+^ responses. While our data indicate a dominant role for TRPV4 channels, contributions from additional ROS-sensitive channels and downstream Ca^2+^ pathways cannot be excluded, consistent with the expression of several ROS-sensitive TRP channels in astrocytes^62–64^. We validated pmFeRIC across *in vitro* and *in vivo* experimental systems, confirming its robustness, reproducibility, and scalability.

A key observation is that astrocytic Ca^2+^ responses differ between *in vitro* and *in vivo* conditions. *In vitro*, RF stimulation elicits a population-wide Ca^2+^ response with prolonged activation, reaching a sustained plateau lasting several minutes. In contrast, *in vivo* astrocytic Ca^2+^ dynamics are substantially faster and are rapidly deactivated, even in the continuous presence of RF stimulation. Moreover, the spatial amplitude (area in µm) of the Ca^2+^ events is an order of magnitude larger *in vivo* with respect to those observed *in vitro*. These differences underscore the importance of studying astrocyte function within intact neural circuits, as simplified experimental models do not faithfully recapitulate physiological cellular interactions.

In visual cortex astrocytes, pmFeRIC evoked Ca^2+^ responses comparable to those induced by locomotion in both amplitude and temporal profile, supporting its efficacy in inducing physiologically relevant Ca^2+^ signaling. Moreover, RF stimulation lasting 1–10 minutes produced Ca^2+^ events with similar amplitude and duration, while longer stimulation increased the proportion of responsive cells. Together, these results indicate that pmFeRIC elicits Ca^2+^ responses constrained by intrinsic astrocytic properties.

Importantly, we show the effects of activating endogenous astrocytic channels on the activity of neighboring neurons. pmFeRIC-mediated astrocytic Ca^2+^ activation increased neuronal spiking and induced both synaptic-dependent and synaptic-independent neuronal Ca^2+^ responses. This effect was presynaptically mediated through an increase in release probability, consistent with previous reports of astrocyte-to-neuron signaling^65,66^. By reproducing established astrocyte–neuron interactions through selective activation of endogenous channels, pmFeRIC supports the evidence that astrocytic modulation of neuronal activity reflects a physiological process rather than an experimental artifact.

Together, these findings establish pmFeRIC as a versatile and physiologically grounded framework for magnetic control of endogenous ion channels. This genetically encoded, cell-specific, and non-invasive approach expands the neuromodulation toolkit and enables causal investigation of astrocyte function, astrocyte–neuron interactions, and glial contributions to neuronal circuitry activity and behavior.

## Lead contact and materials availability

Further information and requests for resources and reagents should be directed to and will be fulfilled by the Lead Contact, Chunlei Liu. Plasmids will be made available upon request and will require a material transfer agreement from C.L. Codes generated here are available at: https://github.com/ChunleiLiuLab/Spiker.

## Acknowledgments

We thank CRL Molecular Imaging Center, UC Berkeley (RRID:SCR_017852) for assistance in confocal microscopy. We thank Emma Stauffenberg for assisting in Ca^2+^ imaging data extraction using the AQuA MATLAB toolbox. We thank Sang Min Han for assisting in writing the MATLAB code for permutation analysis. We thank Professors Sam Pimentel and Dan Feldman for advising in statistical data analysis. Research reported in this publication was supported by the National Institutes of Health (NIH) under awards R01NS129888 and R61AG094651 (C.L.) and U01NS137449 (N.J.), as well as by the Weill Neurohub (N.J.). The content is solely the responsibility of the authors and does not necessarily represent the official views of the NIH.

## Author contributions

C.L. and M.H.-M. conceived, designed, and supervised the project. M.H.-M. designed the experiments; developed all FeRIC constructs; performed *in vitro* and *in vivo* Ca^2+^ imaging, immunoassays in cultured astrocytes, and data analyses; curated the experimental results and wrote the manuscript. J.M.-Z. performed electrophysiological recordings, analyzed electrophysiological data, and developed Python code for neuronal Ca^2+^ spiking and electrophysiological data analyses. T.T. performed *in vitro* Ca^2+^ imaging and immunoassays, analyzed data, and developed code for permutation-based statistical tests to determine cell responsiveness. V.H. designed and built the RF coils and developed the 3D-printed head bar and head holder for *in vivo* experiments. N.T. subcloned FeRIC constructs into AAVs and managed AAV packaging with the resources and supervision of L.T. R.G.N. performed cranial window implantation and visual cortex injections and assisted with *in vivo* Ca^2+^ imaging with the resources and supervision of N.J. Y.L. performed immunoassay labeling in brain slices from *in vivo* experiments. All authors reviewed, edited, and approved the final manuscript.

## Materials and methods

All animal experiments were approved by the UC Berkeley Animal Care and Use Committees and were in accordance with the NIH Guide for the Care and Use of Laboratory Animals and the Public Health Policy.

### Mice

Adult wildtype C57BL/6 mice of both sexes were used for all experiments. Mice were housed in an animal facility with a 12-hour light/dark cycle, an ambient temperature of 20-26°C, and 40-60% humidity.

### Primary neuron cultures

Rat hippocampal neurons were obtained as previously described^1^. Hippocampal tissue from embryonic day 18 Sprague– Dawley rats of either sex (Transnetyx; tissue provided by BrainBits, #SDEHP) was enzymatically treated with a neuron isolation enzyme containing papain (PI88285, Thermo Fisher Scientific Inc.) for 10 min at 37 °C. Tissue was then mechanically dissociated in Minimum Essential Medium (MEM; Transnetyx, tissue by BrainBits). Cells were plated onto poly-D-lysine-coated glass coverslips (1 mg mL^−1^; Sigma, #P0899; 12 mm diameter) in Neurobasal Plus medium (Thermo Fisher Scientific Inc., #A3582901) supplemented with 5% fetal bovine serum (FBS).

For astrocyte-enriched cultures, 2.5 × 10^4^ cells were seeded per coverslip, and 24 h after plating, the culture medium was replaced with Neurobasal medium supplemented with 1% G5 (Thermo Fisher Scientific Inc., #17503012) and 1.5 mM GlutaMAX− (Thermo Fisher Scientific Inc., #35050061). For neuron–astrocyte cocultures, 7.5 × 10^4^ cells were seeded per coverslip, and 24 h after plating, the culture medium was replaced with Neurobasal medium supplemented with 2% B-27 (Thermo Fisher Scientific Inc., #17504044) and 1.5 mM GlutaMAX−.

Cells were maintained at 37 °C in a humidified incubator with 5% CO_2_, and culture medium was refreshed every 3–7 days.

### Plasmids and AAV

The constructs TRPV4^FeRIC^ and TRPV4^FeRIC^ fused to mCherry were obtained as previously described. The mCherry red fluorescent protein encoded in the pLVX-Ef1a-TRPV4^FeRIC^-IRES-mCherry construct was used as a reporter of TRPV4^FeRIC^ expression.

To generate pmFeRIC, restriction sites were introduced into the Lck–mScarlet-I plasmid (pEGFP-C1 backbone, CMV promoter; Addgene, #98821) between the lck membrane-targeting sequence and mScarlet-I, with a 5′ NotI site and a 3′ BamHI site. The FeRIC element corresponding to domain 5 (D5) of human kininogen 1 was subcloned into the NotI and BamHI sites of the pEGFP-C1 backbone vector. Cloning and sequence verification were performed by GenScript Biotech Corporation. To achieve astrocyte-specific expression of pmFeRIC, the lck– D5–mScarlet-I fragment was subcloned into the AAV vector pZac2.1 with the astrocyte-specific GfaABC1D promoter. PCR amplification of lck–D5–mScarlet-I with 5’ EcoRI and 3’ XbaI restriction sites were performed with SuperFi II (Invitrogen 12349050). After restriction digest with EcoRI (NEB R3101) and XbaI (NEB R0145), ligation was performed with T4 DNA ligase (NEB M0202) and transformed with NEB Stable competent E.coli (NEB C3040). After cultures were incubated at 30°C for ∼20 hours, endotoxin-free DNA preps were performed with ZymoPURE II Plasmid Maxiprep Kit (Zymo Research D4202). Whole-plasmid sequence verification was performed by Plasmidsaurus. The resulting construct, pZac2.1-GfaABC1D-lck-D5-mScarlet-I, was packaged into adeno-associated virus serotype 5 (AAV5). AAV particles were packaged and purified either by University of North Carolina Neurotools or prepared in-house. The mScarlet-I red fluorescent protein was used as a reporter of pmFeRIC expression. The constructs GCaMP6m (pGP-CMV-GCaMP6m, Addgene #40754), lck:GFP (Addgene #61099), and ER:GFP (Addgene #182876) were obtained from Addgene.

AAVs encoding GCaMP6 for astrocytic and neuronal expression were obtained from Addgene (AAV5:pZac2.1:GfaABC1D-lck-GCaMP6f, #52924; AAV5:pZac2.1:GfaABC1D-cyto-GCaMP6f, #52925; pAAV:Syn:GCaMP6f.WPRE.SV40, #100837). Constructs driven by generic promoters (Ef1a and CMV) were used in astrocyte-enriched cultures. Constructs driven by cell-type-specific promoters (GfaABC1D and Syn) were used in astrocyte-enriched cultures, neuron–astrocyte cocultures, and *in vivo* transcranial injections.

### Chemical transfection of astrocyte-enriched cultures and neuron-astrocytes cocultures

Astrocyte-enriched cultures were transfected 4-10 days after seeding using Lipofectamine LTX PLUS reagent (Thermo Fisher Scientific Inc., #15338100). Neuron-astrocyte cocultures were transfected 7-8 days after seeding using Lipofectamine− 2000 Transfection Reagent (Thermo Fisher Scientific Inc., #11668027). For all transfections, Opti-MEM reduced-serum medium (Thermo Fisher Scientific Inc., #31985088) was used to prepare DNA-Lipofectamine complexes.

For Lipofectamine LTX–mediated transfection, the transfection mixture per coverslip/well consisted of 100 µL Opti-MEM, 2 µL Lipofectamine LTX, 1.5 µL LTX reagent, 0.3–0.7 µg FeRIC channel DNA, and 0.3–0.5 µg GCaMP6 DNA. Cells were incubated with the transfection mixture at 37 °C for 24 h, after which the medium containing the transfection mixture was replaced with fresh Neurobasal medium supplemented with G5 and GlutaMAX−. Experiments were performed 3-10 days after transfection.

For Lipofectamine 2000–mediated transfection, the transfection mixture per coverslip/well consisted of 100 µL Opti-MEM, 2.5 µL Lipofectamine 2000, 0.3–0.7 µg FeRIC channel DNA, and 0.3–0.5 µg GCaMP6 DNA. Cells were incubated with the transfection mixture at 37 °C for 2-3 h, after which the medium containing the transfection mixture was replaced with fresh Neurobasal medium supplemented with B-27 and GlutaMAX−. Experiments were conducted 5-14 days after transfection.

### Transduction with AAV

Astrocyte-enriched cultures and neuron-astrocyte cocultures were transduced 6–7 days after seeding. All AAV serotypes were diluted to a final concentration of 1 × 10^11^ genome copies (GC) mL^−1^ in Neurobasal medium supplemented with G5 for astrocyte-enriched cultures or Neurobasal medium supplemented with B-27 for neuron-astrocyte cocultures. Cells were incubated with AAV at 37 °C for 48 h, after which the medium was replaced with the corresponding fresh Neurobasal medium. Experiments were performed 5-14 days after transduction.

### RF coil

All RF coils were built as a single loop of copper foil. To enable efficient RF generation at 180 MHz, the RF coils were tuned and matched to ∼50 Ω at 180 MHz with low-loss capacitors (1111C Series, Passive Plus). The capacitor values varied depending on coil size, and the tuning and matching network capacitors were series, then parallel or parallel then series depending on capacitor value availability. The coils were checked to still be matched near 180 MHz when parasitic capacitance due to the nearby metal (e.g., from a microscope) was present.

To generate 180 MHz magnetic fields, the RF signal was generated by a broadband (35 MHz to 4.4 GHz) signal generator (RF Pro Touch, Red Oak Canyon) and amplified using a 5 W amplifier (Amplifier Research, model 5U100, 500 kHz to 1,000 MHz). The magnetic field subsequently produced by the coils was measured using EMC near-field probes (Beehive Electronics model 100B) connected to a spectrum analyzer (RSA3015E-TG, RIGOL Technologies). The Beehive 100B probe was provided with magnetic field calibration data from the manufacturer, and proper scaling factors for 180 MHz measurements were interpolated based on this data. The RIGOL spectrum analyzer was manufacturer-calibrated with a system registered to ISO9001:2008. Ferrite beads designed to block frequencies near 180 MHz (Laird-Signal Integrity Products manufacturer part number 28A0592-0A2) were placed on the cables to prevent them from acting as antennas.

RF coil sizes were chosen based on the mechanical constraints of each experimental setup. For *in vitro* experiments, a 10 mm inner diameter RF coil fabricated on a printed circuit board (PCB) was integrated with the microscope stage. For *in vivo* experiments, a 33.4 mm diameter RF coil built with copper tape on a 3D printed loop was integrated with a 3D printed mouse head holder. This coil was tuned with a series capacitor and matched with a parallel capacitor. The series capacitor was placed at the midpoint of the RF coil loop, saving space and reducing radiation effects.

### AAV delivery and cranial window implantation

Mice were anesthetized with isoflurane (1–3% in O_2_) and administered buprenorphine (0.1 mg kg^−1^, intraperitoneal) and dexamethasone (2.5 mg kg^−1^, subcutaneous). Once fully anesthetized, mice were placed in a stereotaxic frame, lidocaine (3 mg kg^−1^) was injected subcutaneously under the scalp, and eyes were protected with ophthalmic ointment. The scalp overlying the skull was shaved and removed, and the skull surface was cleaned with 2% hydrogen peroxide.

A craniotomy was performed over the primary visual cortex (V1) using stereotaxic coordinates (Bregma −3.0 to −4.5 mm, 2– 3 mm lateral to midline). The skull was thinned and removed (∼2.5 mm^2^) using a dental drill equipped with an FG 1/4 drill bit, exposing the underlying cortex. Bleeding from the dura and wound margins was controlled with sterile, saline-saturated Gelfoam.

For viral delivery, a sterile glass micropipette (15–20 µm tip diameter; Drummond Scientific Company) backfilled with mineral oil and preloaded with AAV vectors was mounted on the stereotaxic arm. Viruses were delivered using a beveled glass pipette (35° angle) with a fitted plunger controlled by a hydraulic manipulator (MO10; Narishige). Multiple injection sites (6-12) were targeted in the left visual cortex. At each site, ∼30 nL of AAV5:gfaABC1D:LCKFeRICmScarlet-I plus AAV5:pZac2.1: gfaABC1D:cyto:GCaMP6f (∼1.8 × 10^13^ GC mL^−1^, ratio 1:1 to 1:3) was injected at ∼250 µm below the pia for Ca^2+^ imaging.

After AAV injection, cranial windows were constructed by bonding a 3 mm or 3.5 mm diameter glass coverslip (#1.5; Zeiss or Fisher Scientific) to a 3D-printed plastic barrel using Norland Optical Adhesive (#61 or #68) and cyanoacrylate, and the assembly was lowered into the craniotomy. Windows were secured to the skull with cyanoacrylate and dental acrylic. A custom-built titanium or polycarbonate head-post was affixed within the surgical site using cyanoacrylate and dental acrylic.

Mice were allowed to recover for ≥ 2 weeks before *in vivo* two-photon imaging. Imaging experiments were conducted in head-fixed anesthetized mice.

### Calcium imaging

Epifluorescence imaging experiments were performed as previously described. Cytosolic Ca^2+^ levels were monitored in cells expressing GCaMP6. Cells expressing FeRIC channels were identified by mCherry or mScarlet-I fluorescence.

Experiments were conducted using an upright AxioExaminer Z.1 microscope (Zeiss) equipped with an Axiocam 702 mono camera controlled by Zen 2.6 software. Excitation light was delivered by an LED unit (33 W cm^−2^; Colibri 7 Type RGB-UV, Zeiss). Fluorescence was recorded with the following filter sets: mCherry or mScarlet-I, excitation 590/27 nm and emission 620/60 nm; GCaMP6, excitation 469/38 nm and emission 525/50 nm. Illumination intensity and exposure times were adjusted to avoid overexposure and minimize GCaMP6 photobleaching. All experiments within a series were performed under identical illumination settings. Images were captured with a W “Plan-Apochromat” 20×/1.0 DIC D = 0.17 M27 75-mm lens at 1 frame s^−1^.

Imaging was performed at room temperature (20–22 °C) using HBSS (Invitrogen, #14025092). Before imaging, cells were washed three times with 1 mL HBSS. Coverslips were then placed in the recording chamber, and cells were initially localized using transmitted bright-field illumination. Fluorescence from mCherry or mScarlet-I and GCaMP6 was confirmed using reflected illumination, and the FOV of interest was selected. In some experiments, a thermocouple coupled to a thermistor readout (TC-344C, Warner Instruments) was placed in direct contact with the HBSS to monitor temperature. Cells were allowed to equilibrate for approximately 20 minutes before imaging.

RF stimulation was delivered using a custom-built RF-emitting coil. Cells were exposed to RF fields at 180 MHz (5–10 µT). Basal Ca^2+^ signaling was recorded in the absence of stimulation for 5–10 minutes prior to RF application.

### Calcium imaging *in vivo* by 2-photon microscopy (2PFM)

*In vivo* 2PFM was performed using a commercial multiphoton microscope (Bergamo® II, Thorlabs). Excitation was provided by a femtosecond Ti:Sapphire laser (Chameleon Ultra II, Coherent), tuned to 920 nm for all experiments. Fluorescence excitation and collection were achieved using a 25× water-immersion objective (Olympus, 1.05 NA, 2 mm working distance). Image acquisition and hardware control were performed using ThorImage software (Thorlabs).

For imaging sessions, mice were anesthetized with isoflurane (1–3% in O_2_) and head-fixed under the objective. Anesthesia was maintained throughout imaging at 0.7–1% isoflurane in O_2_.

Fields of view (FOVs) were selected based on the co-expression of pmFeRIC (mScarlet-I) and GCaMP6. Images were acquired at 3.3 frames/s and a pixel size of 0.753 µm. During analysis, 10 consecutive frames were binned to represent a single time point, leading to an effective frame rate of 0.33 frames/s. Fluorescence signals from mScarlet-I and GCaMP6f were collected sequentially. The RF stimulation coil was embedded within a custom-designed 3D-printed resin head-bar holder; all components of the mouse holder were fabricated from 3D-printed resin to minimize parasitic currents in the RF coil.

Fluorescence imaging was performed during baseline conditions (no stimulation), during RF stimulation (180 MHz, 50 µT), and during a post-stimulation recovery period. Mice were imaged for up to 4 days over a two-week period following recovery from surgery, with one to two unique FOVs acquired per imaging session. In control experiments, mice expressed GCaMP6 alone.

### Electrophysiology: whole-cell patch-clamp recordings

Electrophysiological properties of the cells were examined using the whole-cell patch-clamp technique. Cultures were transferred to a recording chamber filled with extracellular saline solution and positioned on a microscope stage (Axio Examiner Z-1, Zeiss). Neuron-astrocyte cocultures were recorded in Live Cell Imaging Solution (GIBCO A59688DJ) supplemented with 5 mM glucose. Whole-cell recordings were obtained from the soma using patch pipettes with resistances of 3–5 MΩ for neurons and 4–9 MΩ for astrocytes.

In voltage-clamp experiments, the holding potential (V_h_) was set to −60 mV, and pipette capacitive currents were compensated by up to 70%. For all experiments, the internal solution contained (in mM): 130 KMeSO_4_, 10 HEPES-K^+^, 7.6 NaCl, 5 ATP-Mg^2+^, and 0.4 GTP-Na^+^, pH 7.2–7.3. The estimated equilibrium potentials at 20° C were as follows (in mV): 74 Na^+^, −102 K^+^, 124 Ca^2+^, and −75 Cl^−^.

Recordings were performed using a MultiClamp 700A amplifier (Axon Instruments). Data were included only if the seal resistance exceeded 1 GΩ in the cell-attached configuration and if the access resistance (R_a_; ∼15–30 MΩ for neurons, ∼25-50 MΩ for astrocytes) remained stable within 20% of the initial value throughout the experiment. R_a_ was monitored by applying a voltage step (−5 mV, 250 ms) and measuring the peak of the transient, followed by a voltage ramp (0 mV, 500 ms) to calculate the input resistance (R_i_) via a linear fit. The membrane resistance (R_m_) was then calculated using the following formula:

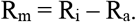

This approach allowed for the simultaneous monitoring of R_a_ and R_m_, ensuring that observed changes in R_m_ were not artifacts of fluctuations in the patch-clamp seal quality.

For neurons recorded in current-clamp mode, no current injection was used to modify the resting membrane potential (I=0). Series resistance (R_s_) was compensated by 80%. R_m_ was monitored throughout the experiment using a current pulse (−20 pA, 250 ms) and an exponential decay fit:

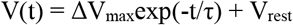

Where the time constant was τ = R_m_C_m_. However, to calculate C_m_ without using R_m_ we used the following equation:

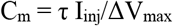

By deriving capacitance from the initial charging trajectory, we decouple the C_m_ estimate from steady-state conductance fluctuations, thereby preventing the propagation of R_m_ estimation errors into the passive membrane calculations.

The current pulse was followed by a ramp (0 pA, 500 ms) to calculate R_m_ using a linear fit.

Data were acquired through a low-pass filter with a cutoff frequency of 3.0 kHz and sampled at 10.0 kHz using an Axon Digidata 1550B A/D converter (Molecular Devices). pClamp 11.2 software (Molecular Devices) was used for data acquisition and the generation of current pulses and voltage steps.

### Cultured cells immunofluorescence staining

Cells were fixed at DIV 11–15 with ice-cold 4% paraformaldehyde (PFA) for 15 min and washed three times with phosphate-buffered saline (PBS) for 15 min. Cells were then treated with blocking solution (1X, Invitrogen, #R37627) according to the manufacturer’s instructions.

For labeling TRP channels at the extracellular domain, cells were not permeabilized and were incubated with primary rabbit anti-TRP antibodies (1:200, Alomone Labs) diluted in 1% bovine serum albumin (BSA) in PBS overnight at 4 °C in a humidity chamber. Samples were washed with PBS and incubated for 1 h at room temperature (RT) with Alexa Fluor 594-conjugated secondary antibody (1:1000, Jackson ImmunoResearch Labs) in 5% BSA in PBS.

For intracellular protein staining (GFAP and MAP2), cells were permeabilized with 0.25% Triton X-100 in 5% BSA for 15 min at RT. For co-immunostaining with TRP channels, permeabilization was performed after the extracellular TRP staining was completed. Cells were then treated with blocking solution (1X, Invitrogen) before primary antibody incubation. Primary antibodies included mouse anti-GFAP (1:500, Cell Signaling Technology, GA5) and chicken anti-MAP2 (1:500, EnCor Biotechnology, polyclonal) and were incubated overnight at 4 °C in a humidity chamber. Following three PBS washes, samples were incubated for 1 h at RT with secondary antibodies Alexa Fluor 647 goat anti-mouse and Alexa Fluor 488 donkey anti-chicken (1:1000 each, Invitrogen) in 5% BSA in PBS.

Nuclei were counterstained with NucBlue Fixed Cell Stain ReadyProbes reagent (Invitrogen, #R37606) according to the manufacturer’s instructions. Samples were mounted on coverslips with Fluoromount-G (SouthernBiotech, #0100-01) and imaged using a confocal microscope (Zeiss LSM 980 NLO).

### Mouse brain tissue immunofluorescence staining

Mice were deeply anesthetized with ketamine (100 mg kg^−1^) and xylazine (10 mg kg^−1^) and transcardially perfused with 50 mL ice-cold phosphate-buffered saline (PBS), followed by 50 mL 4% paraformaldehyde (PFA) in PBS. Brains were removed, post-fixed in 4% PFA at 4 °C for 2-4 h, and cryoprotected sequentially in 20% and then 30% sucrose in 0.1 M PBS at 4 °C until they sank. Coronal sections (20 µm) were cut from caudal to rostral using a cryostat (Microm HM 525, Thermo Fisher Scientific Inc.).

Sections were washed in PBS, permeabilized with 0.3% Triton X-100 in PBS (0.3% PBST) for 30 min at room temperature (RT), and blocked for 90 min at RT in 0.1% PBST containing 5% normal donkey serum and 1% bovine serum albumin (BSA). Sections were incubated with primary antibodies overnight at 4 °C, followed by incubation with secondary antibodies for 1 h at RT. Nuclei were counterstained with NucBlue Fixed Cell Stain ReadyProbes reagent (Invitrogen, #R37606) according to the manufacturer’s instructions, and sections were mounted in antifade medium. Images were acquired using a laser-scanning confocal microscope (ZEISS LSM 900).

Primary antibodies: anti-GFAP (1:500, Cell Signaling Technology, #3670) and anti-mScarlet-I (1:500, Bioss, #bs-33856R). Secondary antibodies: goat anti-mouse IgG (H+L) Alexa Fluor 488 (1:1000, Invitrogen, #A-11001) and donkey anti-rabbit IgG (H+L) Alexa Fluor 555 (1:1000, Invitrogen, #A-31572).

## Data analysis

### Calcium imaging analysis – Detection of calcium events using the MATLAB AQuA algorithm

For *in vitro* Ca^2+^ imaging, raw imaging data were converted from .czi to TIFF format using Fiji (ImageJ; NIH). Background fluorescence was subtracted using a rolling-ball algorithm (radius, 50 pixels), and images were spatially filtered with a 2-pixel median filter to reduce noise. For *in vivo* Ca^2+^ imaging, raw data were saved directly as TIFF files, and no preprocessing was applied prior to analysis.

Astrocytic Ca^2+^ signaling was analyzed using the MATLAB-based AQuA algorithm to detect Ca^2+^ events and quantify their amplitude and spatiotemporal properties. For *in vitro* datasets, the GCaMPExVivo pipeline was used, whereas the GCaMP6InVivo pipeline was applied to *in vivo* datasets. Analysis parameters were held constant across all datasets and included an intensity threshold of 5, a smoothing factor of 1, a minimum event size corresponding to ∼6 µm, and a z-score threshold of 3.

Following event extraction, data visualization and statistical analyses were performed using OriginLab software. For population-level comparisons, Ca^2+^ events from all cells within a single field of view were pooled and compared across experimental conditions. For single-cell responsiveness analyses, Ca^2+^ events detected in individual cells were analyzed separately using a permutation-based statistical test.

### Peak-scaled Non-Stationary Fluctuation Analysis (psNSFA)

Miniature excitatory postsynaptic currents (mEPSCs) were selected based on their rise time (e.g., < 2 ms) to include events occurring closer to the soma and minimize the filtering effects of dendritic axial resistance. These mEPSCs were aligned by their maximum rising slope, and an average template was calculated. The average response was then scaled to the peak of each individual event. To account for noise, the peak amplitude of each mEPSC was determined by averaging the signal within a 400 µs window centered on the peak position.

The scaled average response was subtracted from each individual mEPSC, yielding fluctuations around the scaled mean. The variance of these fluctuations was calculated and plotted against the binned amplitude (1 pA bins) of the average response.

The resulting relationship was fit using the following equation:

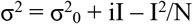

Where:

- σ^2^ is the variance around the scaled mean.
- σ^2^0 is the basal (background) variance of the recording.
- i is the single-channel (unitary) current.
- I is the average binned current.
- N is the average number of functional channels.

### Automated event detection and signal processing (Spiker, Python analysis)

To detect and analyze events in electrophysiological and Ca^2+^ imaging recordings, custom Python-based software was employed. Configurations were tailored to each recording type based on the acquisition rate, noise floor, and the specific kinetics of the events to be detected, such as mEPSCs, action potentials (APs), or Ca^2+^ spikes.

#### Noise Estimation and Thresholding

Each recording was segmented into intervals smaller than the typical inter-event interval (resp_increment, ranging from milliseconds to seconds) to identify segments devoid of events. The standard deviation (SD) of the noise within these segments was calculated to construct the std_resp vector. This vector was subsequently segmented into larger intervals (std_increment, ranging from seconds to minutes), and the minimum SD values within each increment were selected to construct the std_min vector. This std_min was then multiplied by a scaling factor (n_deviations) and interpolated across the entire recording.

A copy of each recording was smoothed using a Gaussian convolution kernel to generate a smoothed_resp vector. The peak_noise vector, which served as the amplitude threshold (AT), was defined as the sum of smoothed_resp and the interpolated std_min. Signal segments exceeding the peak_noise were identified as putative events.

#### Derivative Analysis and Event Selection

The first derivative of the signal was subjected to the same procedure; slopes exceeding their corresponding peak_noise were classified as the slopes of putative events. Generally, events preceded by a high-magnitude slope in the direction of the event were further refined based on the specific event type.

For brevity, the key selection parameters included:

- slope_peak_time
- peak_to_peak
- max_rise_time
- min_amplitude
- max_slope
- min_auc (Area Under the Curve)

### Statistical analysis

All experimental groups included at least three independent experiments. Statistical analyses were performed using OriginPro 2021 (v.9.8; OriginLab). Data are presented as mean ± s.e.m.

Normality was assessed for each dataset. For normally distributed data, comparisons between two groups were performed using a two-sample Student’s t test, and comparisons among three or more groups were performed using one-way ANOVA followed by Bonferroni’s multiple-comparisons test. For non-normally distributed data, comparisons between two groups were performed using the two-sample Kolmogorov– Smirnov test, and comparisons among three or more groups were performed using one-way Kruskal–Wallis ANOVA followed by Dunn’s multiple-comparisons test. Statistical significance was defined as p < 0.05 (*), p < 0.001 (**), or p < 0.0001 (***).

### Permutation test for cell responsiveness

Ca^2+^ event parameters were extracted for each cell using the MATLAB toolbox AQuA. Cell responsiveness to RF stimulation was assessed using a permutation test based on the area under the curve (AUC) and ΔF/F_0_. A custom MATLAB script was applied to all Ca^2+^ events recorded from individual astrocytes co-expressing FeRIC and GCaMP to test for a significant increase in Ca^2+^ activity following RF stimulation. Event parameters obtained under basal (no RF) and RF conditions were concatenated and randomly reassigned into two groups (pseudo-basal and pseudo-RF). The mean of each group was calculated, and the difference between means was computed. This procedure was repeated 10,000 times per cell to generate a null distribution. The observed difference between the basal and RF conditions was then compared against the null distribution to determine RF responsiveness at the single-cell level. Cells were classified as responsive if they exhibited a significant increase in Ca^2+^ activity (one-tailed test, p < 0.05). Cells not meeting this criterion were classified as non-responsive.

## Extended Data

**Extended Data Fig. S1.**
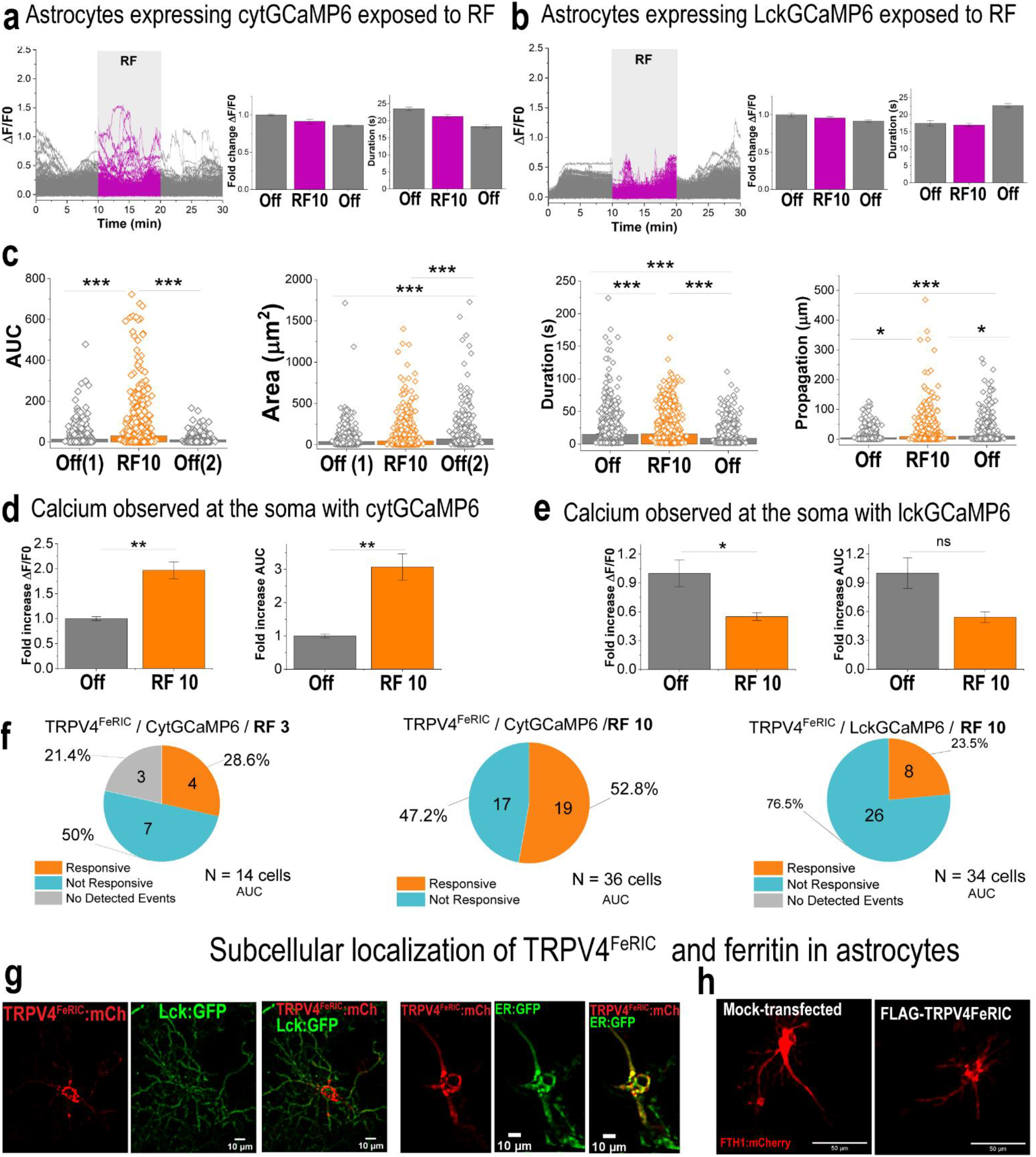
Effects of RF stimulation on Ca^2+^ activity in control astrocytes expressing GCaMP6 alone and in astrocytes expressing TRPV4^FeRIC^, in which the channels are predominantly localized to the endoplasmic reticulum. (**a**,**b**) Fluorescence traces from astrocytes expressing cytGCaMP6 (n= 4 independent experiments, 32 cells, 2020 – 4302 Ca^2+^ events; a) or lckGCaMP6 (n = 4 independent experiments, 19 cells, 684 -2099 Ca^2+^ events; b) during alternating 10-min RF off/on periods. (**c-e**) Mean (± SEM) cytGCaMP6 AUC, area, duration, and propagation of Ca^2+^ events in TRPV4^FeRIC^–expressing astrocytes during 10-min RF stimulation, quantified in astrocytic processes or soma using cytGCaMP6 (c,d) or lckGCaMP6 (e). (**f**) Percentage of astrocytes responsive to 3-or 10-min RF stimulation when expressing TRPV4^FeRIC^ and cytGCaMP6 (left, middle) or lckGCaMP6 (right). (**g**,**h**) Live imaging of astrocytes expressing TRPV4^FeRIC^:mCherry together with lck:GFP (plasma membrane marker; g) or ER:GFP (endoplasmic reticulum marker; h). Cell responsiveness was determined using the permutation test for the difference in means. Where applicable, *P < 0.05, **P < 0.001, or ***P < 0.0001

**Extended Data Fig. S2.**
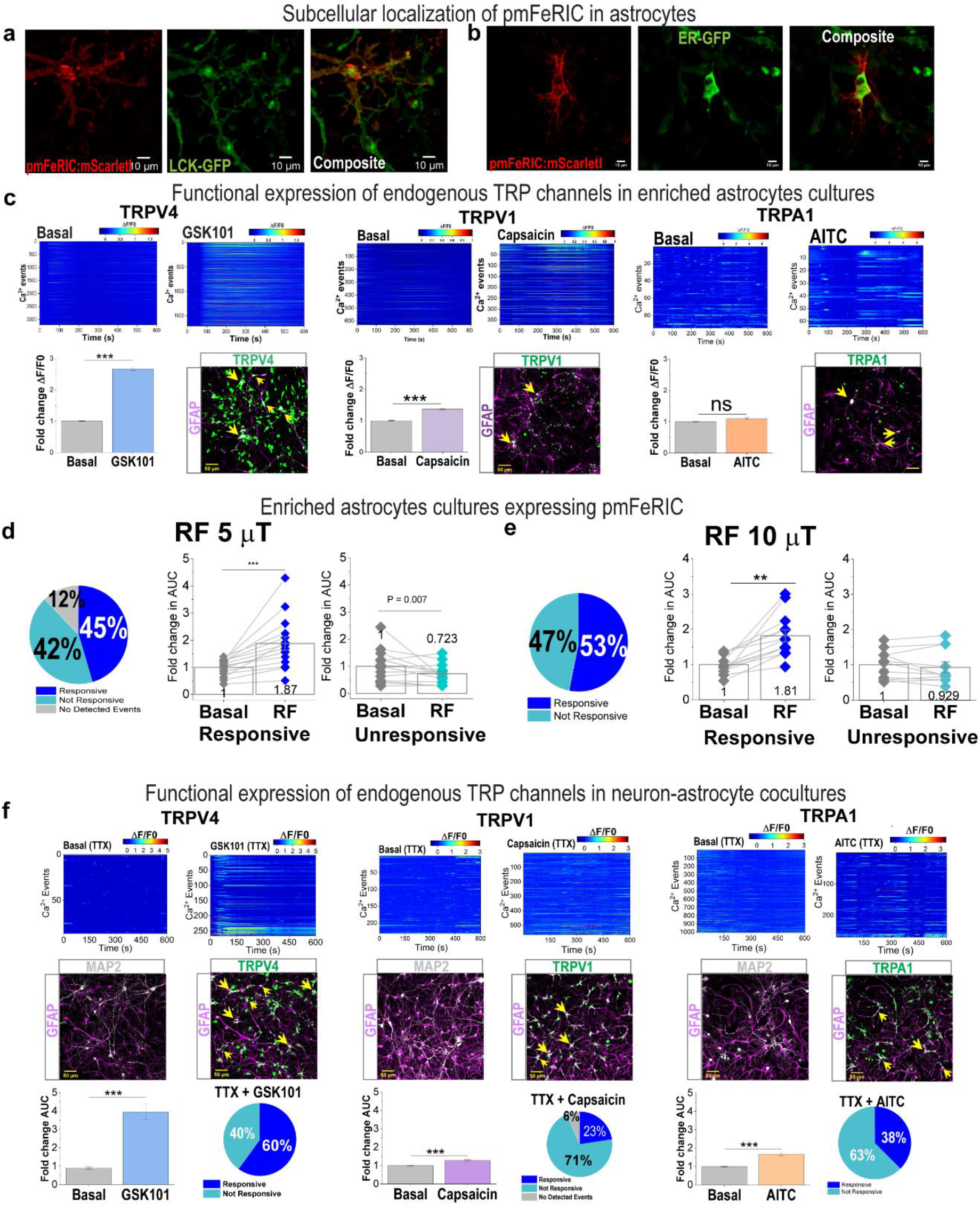
Subcellular localization of pmFeRIC and expression of endogenous TRP channels in astrocytes. (a,b) Live imaging of astrocytes coexpressing pmFeRIC:mScarletI and lckGFP (plasma membrane marker; a) or ER–GFP (endoplasmic reticulum marker; b). (c) Heat maps and fold change in Ca^2+^ events amplitude (ΔF/F0) evoked by selective agonists of TRPV4, TRPV1, and TRPA1 in astrocytes from enriched astrocyte cultures. Channel expression was confirmed by immunolabeling for TRPV4, TRPV1, and TRPA1 (green). Yellow arrows indicate cells coexpressing GFAP and TRP channels. (d,e) Percentage of RF-responsive pmFeRIC-astrocytes following 10-min stimulation at 5 µT (d) or 10 µT (e). Fold change in AUC is shown separately for responsive and nonresponsive cells. (f) Heat maps and fold change in Ca^2+^ events amplitude (ΔF/F0) induced by selective agonists of TRPV4, TRPV1, and TRPA1 in astrocytes from neuron– astrocyte cocultures. Channel expression was confirmed by immunolabeling for TRP channels (green), together with GFAP and MAP2. Yellow arrows indicate cells coexpressing GFAP and TRP channels. Bottom panels show fold change in AUC and the percentage of cells responsive to each TRP channel agonist.

**Extended Data Fig. S4.**
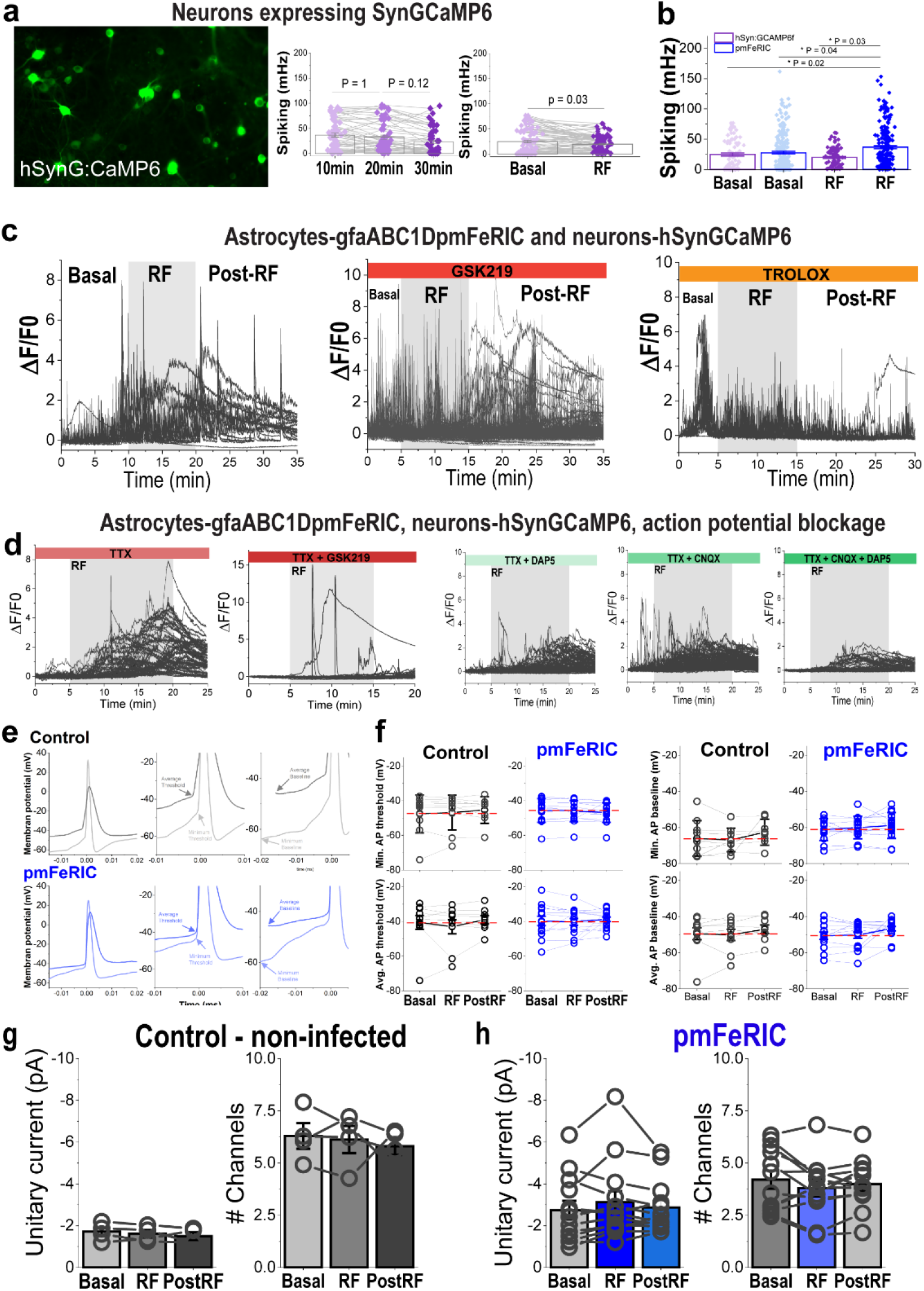
Pharmacological and electrophysiological characterization of pmFeRIC astrocyte–mediated control of neuronal activity. (**a**) Control neurons expressing hSyn:GCaMP6f imaged for up to 30 min without stimulation (n = 3 independent experiments, 55 cells) or during RF stimulation (10 µT; 5 independent experiments, 88 cells). Neuronal Ca^2+^ spiking frequency is shown. (**b**) Comparison of neuronal Ca^2+^ spiking frequency in cocultures without (GCaMP6f only) or with pmFeRIC-expressing astrocytes under basal conditions and following RF stimulation. (**c**,**d**) hSyn:GCaMP6f fluorescence traces from neurons cocultured with pmFeRIC-expressing astrocytes during basal, RF, and post-RF periods in the absence or presence of TTX and pharmacological blockers, including the TRPV4 antagonist GSK219, the ROS scavenger Trolox, and ionotropic glutamate receptor antagonists. (**e**,**f**) Representative action potential waveforms and summary data (mean ± SEM) from neurons cocultured without (control; n = 80 to 10 cells) or with pmFeRIC-expressing astrocytes (n = 13 to 16 cells). Traces show full action potential profiles (left), highlighting minimum and average action potential thresholds (middle) and baseline membrane potential parameters. (**g**) Estimated unitary current and inferred number of active channels during basal, RF, and post-RF conditions from neurons cocultured without (control; n = 4 cells) or with pmFeRIC-expressing astrocytes (n = 13 cells). For Ca^2+^ imaging data, statistical significance was assessed as indicated in Fig.1. For patch-clamp data, statistical significance was assessed using a parametric one-way ANOVA with Bonferroni’s post hoc test; P values are indicated.

**Extended Data Fig. S5.**
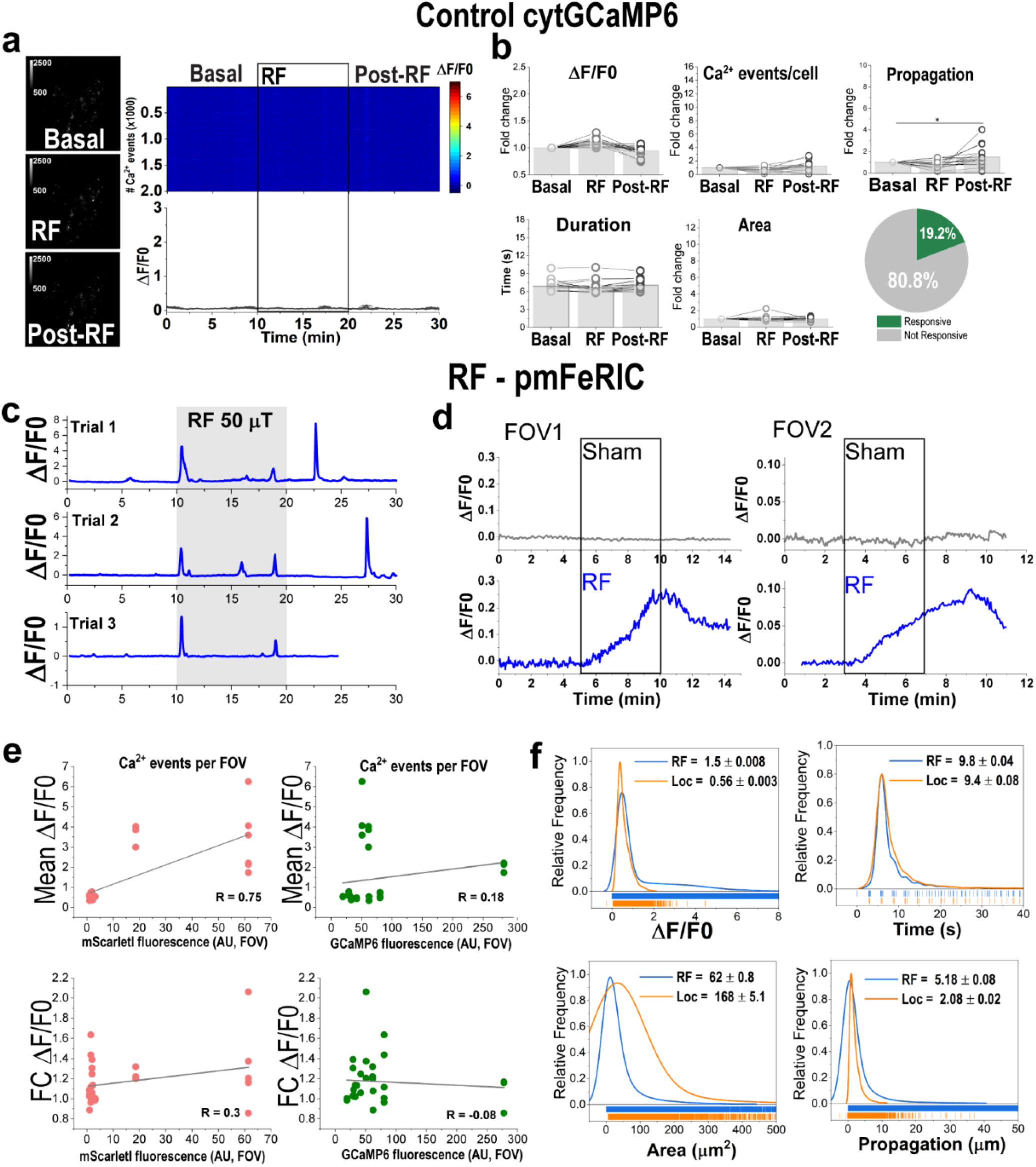
Magnetically induced Ca^2+^ responses in V1 astrocytes. (**a**) cytGCaMP6 fluorescence in V1 astrocytes expressing only cytGCaMP6 during basal, sham, and post-RF periods. Representative heat map of Ca^2+^ events and averaged cytGCaMP6 fluorescence traces from all detected events observed in an independent experiment. (**b**) Mean (± SEM) Ca^2+^ event properties measured during basal, RF, and post-RF periods in control astrocytes (4 mice, 11 independent experiments, 244 cells). (**c**) Mean cytGCaMP6 fluorescence changes across a single field of view during three consecutive 10-min RF stimulations, separated by 20-min rest periods, in the same pmFeRIC-expressing mouse. (**d**) Mean cytGCaMP6 fluorescence changes across two fields of view in the same pmFeRIC-expressing mouse during either sham or 5 min RF stimulation. (**e**) Relationship between mean maximum ΔF/F0 and mScarlet-I fluorescence (left) or cytGCaMP6 fluorescence (right). (**f**) Distribution of Ca^2+^ events properties during RF stimulation compared with those observed under locomotion activity. Statistical analysis was performed as described in Fig. 1.

